# SCO-pH: Microfluidic dynamic phenotyping platform for high-throughput screening of single cell acidification

**DOI:** 10.1101/2024.05.08.593179

**Authors:** Hyejoong Jeong, Emilia A. Leyes Porello, Jean Gabriel Rosario, Da Kuang, Syung Hun Han, Jai-Yoon Sul, Bomyi Lim, Daeyeon Lee, Junhyong Kim

**Affiliations:** Department of Chemical and Biomolecular Engineering, University of Pennsylvania, Philadelphia, PA 19104, USA; Department of Biology, University of Pennsylvania, Philadelphia, PA 19104, USA; Department of Computer and Information Science, University of Pennsylvania, Philadelphia, PA 19104, USA; Department of Bioengineering, University of Pennsylvania, Philadelphia, PA 19104, United States; Department of Systems Pharmacology and Translational Therapeutics, Perelman School of Medicine, University of Pennsylvania, Philadelphia, PA 19104, USA

**Keywords:** single cell, microfluidics, extracellular acidification, high-throughput, phenotyping, glycolysis

## Abstract

Studies on the dynamics of single cell phenotyping have been hampered by the lack of quantitative high-throughput metabolism assays. Extracellular acidification, a prominent phenotype, yields significant insights into cellular metabolism, including tumorigenicity. Here, we develop a versatile microfluidic system for single cell optical pH analysis (SCO-pH), which compartmentalizes single cells in 140-pL droplets and immobilizes approximately 40,000 droplets in a two-dimensional array for temporal extracellular pH analysis. SCO-pH distinguishes cells undergoing hyperglycolysis induced by oligomycin A from untreated cells by monitoring their extracellular acidification. To facilitate pH sensing in each droplet, we encapsulate a cell-impermeable pH probe whose fluorescence intensities are quantified. Using this approach, we can differentiate hyperglycolytic cells and concurrently observe single cell heterogeneity in extracellular acidification dynamics. This high-throughput system will be useful in applications that require dynamic phenotyping of single cells with significant heterogeneity.

## 1. Introduction

Cellular acidification is a key indicator of mitochondrial respiration and glycolysis, giving an insight into cellular metabolic function. Typically, cancer cells are characterized by high glucose metabolism and lactate production through aerobic glycolysis, also known as the Warburg effect.[1] Hence, the detection of cellular acidification is of great value in basic cell biology as well as for diagnosis and therapeutic monitoring. Recently, the rise of single cell biology has shown that even for cells of the same genotype and same differentiated identity, there are individual variations leading to broad heterogeneity in many traits including metabolic functions such as acidification.[2–7] It is essential to quantify this heterogeneity because of its role in determining drug sensitivity, cellular response, and phenotype.

Fluorescence-activated cell sorting (FACS) is a widely used tool for single cell phenotyping. FACS typically detects and sorts cells by their surface or intracellular fluorescent markers. However, this method is not suitable for secretion analysis, and is only a snapshot that does not contain dynamics over a period. Moreover, cells are fixed and no longer alive after FACS, limiting its utility. For dynamic analysis of cellular metabolism, Agilent Seahorse XF Analyzer is commonly used as a practical tool. It simultaneously measures extracellular acidification rates and oxygen consumption rates of live cells. Also, this instrument has the advantage of measuring cellular response in real-time upon addition of stimulants during cell culture and being a label-free method. However, Seahorse is limited to bulk cell measurement and does not allow for single cell analyses. [8–10]

Droplet-based microfluidic devices have emerged as a powerful high-throughput platform for single cell analysis because of their ability to allow compartmentalization of single cells into tiny chambers and analyses of metabolic products[11], proteins[12–14], nucleic acids[15–17], phospholipids[18] and cellular mechanics[19]. This technology has been successfully commercialized for single cell transcriptomics library for next generation sequencing.[20] Single cell phenotyping can be achieved through observation of individual cells and their intracellular readouts. Moreover, droplet-based methods allow isolation of individual cells in micro chambers, which in turn allows the detection of cells’ metabolic outputs into their surrounding media. For example, droplet-based single cell immunoglobulin (IgG) quantification and secretion kinetics have been reported.[13] This “DropMap” is a high-throughput system that allows analysis of IgG release in 40,000 droplets at a time. This method is based on paramagnetic nanoparticles and antibodies for quantitative detection of IgG, but its effectiveness is constrained by the range of IgG detection, the quality of nanoparticles, and the diverse characteristics of antibodies. Another study has reported droplet-based lactate quantification method from single cells by using a commercial lactate assay kit. However, this method relies on a complicated enzymatic reaction to detect lactate and only allows for measurements up to tens of minutes, thus limiting its utility in long-term monitoring of cell metabolism.[11]

Herein, we develop a novel technology, SCO-pH (single cell optical pH analysis microfluidic platform), based on our years of research experience on microfluidic technologies.[21–28] It is a high-throughput single cell acidification analysis platform that co-encapsulates single cells and a cell-impermeant ratiometric pH probe, SNARF™-4F 5-(and-6)-Carboxylic Acid. 40,000 droplets are arranged into a microwell array and monitored for at least 3 hours in parallel. This technology offers several advantages: First, cells are encapsulated in hermetic droplets, and we measure extracellular pH, a cellular property independent of the measurement technique, making it a highly biologically relevant marker.[11] This technique is quantitative, allowing measurements of global acidification thus enabling tracking of glycolysis and possibly other cellular metabolisms. Second, 10’s of thousands of single cell droplets are measured in parallel by positioning droplets in a microwell array. Lastly, this technique is physically and biologically straightforward to set up and conduct. The technique uses a commercial pH probe that is robust and non-toxic. Other dyes can potentially be added for multiplexed analyses. Building upon our prior success in selectively releasing droplets from a microwell array,[29] we expand the functionality of the microwell array to enable pH quantification and automated data analysis, providing a powerful and versatile tool for cellular metabolic phenotyping and clinical cellular diagnosis.

## 2. Materials and methods

### 2.1. Reagents

Fluorinert FC-40, and trichloro(1H,1H,2H,2H-perfluorooctyl)-silane (PFOTS), and oligomycin A were purchased from Sigma-Aldrich (MO, USA). HFE-7500 was obtained from 3M (MN, USA). FluoSurf 2wt% in HFE-7500 was obtained from Dolomite Microfluidics (Royston, UK). Sylgard™ 184 silicone elastomer kit was purchased from Dow Corning (MI, USA). Invitrogen™ SNARF™-4F 5-(and-6)-Carboxylic Acid, CellTrace^TM^ Violet, Calcein AM, and live cell imaging solution were purchased from Invitrogen (MA, USA). Seahorse XF RPMI assay medium pack is purchased from Agilent Technologies (CA, USA).

### 2.2. Device fabrication

Devices for droplet generation (called “droplet generator”) and trapping droplets (called “microwell array”) were fabricated using a previously reported method.[29] The device masks were designed by AutoCAD 2018 and printed on Cr masks by CAD/Art Service, Inc. (CA, USA). A master mold was fabricated on a 3″ silicon wafer (University Wafer Inc., MA, USA) using the conventional soft lithography technique. The master molds were all fabricated inside a cleanroom in the Quattrone Nanofabrication Center of the Singh Center of Nanotechnology at the University of Pennsylvania. The droplet generator was fabricated via the single-layer soft lithography technique. A negative photoresist SU-2025 (MicroChem, MA, USA) and a film mask was used, and the thickness of the droplet generator mold was 40 μm controlled by adjusting the rotation speed of spin coating in conjunction with the UV exposure time under a mask aligner (ABM 3000HR Mask Aligner, ABM, NY, USA). To produce the master for the bottom trap channel of a microwell array, a multilayer mold fabrication method was employed using Cr photomasks prepared by a laser writer (Heidelberg DWL 66+ Laser Writer, Heidelberg Instruments, Germany). Multilayer mold fabrication skipped mold development after initial post-bake and proceeded with spin coating of the second photoresist layer. The top flow channel mold was fabricated via the single-layer soft lithography technique. Fabricated master molds were subsequently silanized with PFOTS to facilitate the detachment of cured polydimethylsiloxane (PDMS). PDMS precursor was prepared by mixing the base and curing agents of Sylgard 184 in a 10:1 ratio and is degassed in a vacuum chamber for 30 min.

The degassed PDMS mixture was poured onto the master mold. The thickness of poured PDMS for a droplet generator and a microwell array was ∼3 mm. After additional degassing step for 30 min, PDMS was cured in an oven for 4 h at 65 °C. The PDMS molds were bonded to a plain glass slide using a conventional oxygen plasma treatment.

### 2.3. Droplet generation and trapping

The droplet generator made with PDMS was silanized with a 2% PFOTS solution in HFE-7500 oil for 5 min following plasma treatment. The device was flushed with neat HFE-7500 oil, then connected with three polytetrafluoroethylene (PTFE) tubing lines, each linked to a syringe. One syringe, filled with HFE-7500 containing 2 wt% EA-surfactant, was connected to an oil inlet at the top of the generator. The other two syringes, filled with an aqueous solution containing cells or SNARF-4F dye, were connected to water inlets in the middle of the device. The EA surfactant is a commercially available tri-block copolymer composed of perfluoropolyether (PFPE) and polyethylene glycol (PEG) blocks and is most widely used in emulsion stabilization in fluorinated oils. By connecting syringes to two syringe pumps (Harvard Apparatus, MA, USA) for each oil and aqueous solution, flow rates of 150 μL/h and 300 μL/h were used for the oil and aqueous phases, respectively. In this condition, uniform droplets with approximately 65 μm in diameter were generated. Prior to droplet introduction, a microwell array was silanized with a method same as above and flushed with neat HFE-7500 oil for 2 min to remove air bubbles within the device. Droplets generated from the droplet generator traveled through a PTFE tubing and immediately entered a microwell array device. With >60% of the wells filled with droplets, the droplet injection tubing was disconnected, and HFE-7500 containing 1 wt% of EA-surfactant and 1 wt% of fluorinated SiO_2_ NPs was injected slowly (200 μL/h) to fill the remaining wells with the droplets (fluorinated SiO_2_ NPs were synthesized in the lab. The protocol and results are provided in the Supporting Information.). To remove untrapped droplets from the channel, neat FC-40 oil was injected rapidly (500 μL/h).

### 2.4. Droplet crosstalk test

Two different ensembles of droplets were generated using droplet generators: one filled with only Seahorse XF RPMI medium supplemented with 1 mM of pyruvate, 2 mM of glutamine, and 10 mM of glucose, and the other filled with 100 μM of SNARF-4F dye in the same buffer. Droplets traveled through a PTFE tubing and were collected in a microtube. The droplet mixture was well mixed while maintaining their stability and injected into a PDMS observation chamber with 40 μm height and 1X1 cm^2^ dimension by using a pipette. Mixed droplets arranged in a single layer for ease of observation were monitored for 3 h under the bright field microscopy and through 600 nm, and 667 nm wavelength fluorescence microscopy.

### 2.5. Cell culture

EL4 (TIB-39) mouse lymphoma cell line was purchased from American Type Cell Culture (VA, USA). The thawed cell pellet was suspended in RPMI 1640 supplemented with 10% v/v fetal bovine serum and 1%(v/v) penicillin-streptomycin. The cell suspension was seeded in a Corning T-75 flask at density of 1.5X10^5^ cells/mL with total volume of 30 mL and cultured at 37°C and 5% CO_2_ in a humidified incubator, with the medium changed every 2-3 days. The cell suspension was collected by centrifugation at 200 rcf for 10 min and the supernatant was removed. The cell pellet was resuspended in a live cell imaging solution or Seahorse XF RPMI assay medium to a density of 2.4X10^7^ cells/mL for cell encapsulation experiments.

### 2.6. Cell encapsulation and trapping

Cell suspension with a density of 2.4X10^7^ cells/mL was prepared into two, and the two groups of cells were stained with Calcein AM and CellTrace^TM^ Violet, respectively. The number of cells was dictated to achieve approximately 30% of the single cell containing droplet generation following a Poisson distribution shown in equation (1).

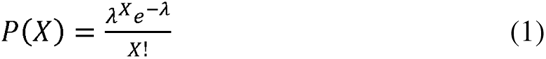

P is probability for the number of cells in a droplet. X is the number of cells in a droplet., 1 is the mean number of cells per droplet and is calculated by multiplying cell concentration by the droplet volume.[30]

For the encapsulated cells, glucose supplemented XF medium was used to detect extracellular pH changes during imaging. Cell encapsulation was performed twice to generate two different droplets: one containing 10 μM of oligomycin for hyperglycolytic cells, and the other without oligomycin serving as a control (untreated cells). To prepare hyperglycolytic cell model, Calcein AM-stained cells were encapsulated by using two aqueous solutions, one filled with 100 μM of SNARF-4F dye and 20 μM of oligomycin, and the other one filled with 2.4X10^7^ cells/mL. For untreated cells, an equal amount of dimethyl sulfoxide (DMSO), which is a solvent for oligomycin stock solution, was added to the SNARF-4F-containing solution. According to the method 2.3., uniform droplets were generated and collected in a microtube. The droplet mixture was well mixed and injected into a microwell array by using a syringe and a syringe pump to maintain the droplets stably with a slow flow rate. The subsequent process was carried out according to the method 2.3.

### 2.7. Time-lapse microscopy imaging

Imaging started immediately after the droplets were captured in the microwell array and was performed using a Zeiss Observer 7 inverted microscope with a motorized stage. The microwell array was located in an environmental chamber that maintained cell culture environment with 95% humidity, 37°C, and 0% of CO_2_ (Okolab, Pozzuoli, Italy). Excitation light was provided by a LED source (Colibri 5, Carl Zeiss, Germany). Fluorescence for the specific channels was recorded using a multifilter (385, 469, 555, and 631 nm) appropriate band-pass filters (SNARF-4F-Cy3: Emission at 600 nm, bandwidth 32 nm; SNARF-4F-Cy5: Emission at 667 nm, bandwidth 30 nm; Chroma Technology Corp. USA) and camera settings (Andor ZL41 Cell 4.2 sCMOS, Oxford Instruments, UK) at room temperature (25°C) and ambient oxygen concentration with 0% of CO_2_. Images were acquired using a 10X objective (Plan-Apochromat, NA 0.45, Zeiss, Germany). To verify the technology, three positions out of 240 positions of an array were acquired every 5 min in three channels over 3 hours. One position in the array was imaged in three channels, and the position was changed afterwards and imaged repeatedly. For the 240 position imaging, we used a custom journal of MetaMorph software to obtain images from an entire device automatically by imaging loops. 240 positions in the microwell array were imaged in three channels in a loop of imaging, and imaging was repeated for 3 hours. The temporal resolution of the imaging is 14 min.

### 2.8. Image analysis

Prior to image analysis, all images were sorted by a custom Python script based on the number of positions. The images were then analyzed using a Fiji free software with Mosaic Particle Tracker plugin and a custom MATLAB script (Mathworks). The details of the parameters used in the Mosaic Particle Tracker were: radius=10, cutoff=0, per/abs=0.6, link=4, displacement=5, and dynamics=“Brownian”. Encapsulated cells in the droplets were detected in Green and UV images using a custom Fiji script for automated and quantitative single- and multiple cells (n>1) tracking.[31] For each position, the custom MATLAB script then generated a binarized droplet mask for 37 images over 3 hours and mapped each cell from the Particle Tracker into the droplets based on the tracked cell x and y coordinate position. The script additionally extracted the fluorescence intensities of the tracked cells in single cell droplets in Green/UV, SNARF-4F-Cy3, and SNARF-4F-Cy5 channels. The Green or UV signals were normalized to remove the bleaching of dyes. Fluorescent intensities of empty droplets were sorted out separately. The mean fluorescence intensities of droplets collected from SNARF-4F-Cy3 and SNARF-4F Cy5 images were used to calculate a pH value following the manufacturer’s protocol.[32]

### 2.9. Data availability

The scripts used in the paper are freely available on GitHub for file organization (https://github.com/kimpenn/microscope-utility) and image analysis (https://github.com/HyejoongJeong/UPenn_pH).

## 3. Results and Discussion

### 3.1. Droplet-based single cell extracellular pH sensing

Single cell encapsulation is a powerful phenotyping approach as metabolites accumulate in picoliter hermetic droplets at sufficient concentrations, allowing their detection at the single cell level. The general principle and workflow of SCO-pH is schematically illustrated in **Fig. 1a**. Single cells are compartmentalized in monodisperse aqueous droplets with 65 μm diameter (∼ 143 pL in volume) in an inert fluorinated oil using a microfluidic flow-focusing device (**Fig. 1a** and **Fig. S2**). Reagents and cells are injected into the microfluidic flow-focusing device, with the cell concentration adjusted to generate up to 30% of droplets containing single cells, following the Poisson distribution; 18% and 52% of droplets are empty and contain multiple cells, respectively. The generated droplets subsequently flow into a device with a serpentine channel, where the top surface features a two-dimensional microwell array. As they flow through the channel, droplets float into the microwells due to the density difference between water (ρ=1.0) and oil (ρ=1.614) (**Fig. S3c**). Approximately 40,000 droplets (i.e. approximately 12,000 single cell droplets) are captured in the microwell array within 3 minutes. A 10× objective lens can capture 13 x 13 microwells in a single imaging window; each well in each image is indexed for kinetic analysis of acidification. An automated microscope stage is controlled to image the entire microwell array device across 240 positions, capturing images of all 40,000 microwells within 15 minutes.

**Figure 1.**
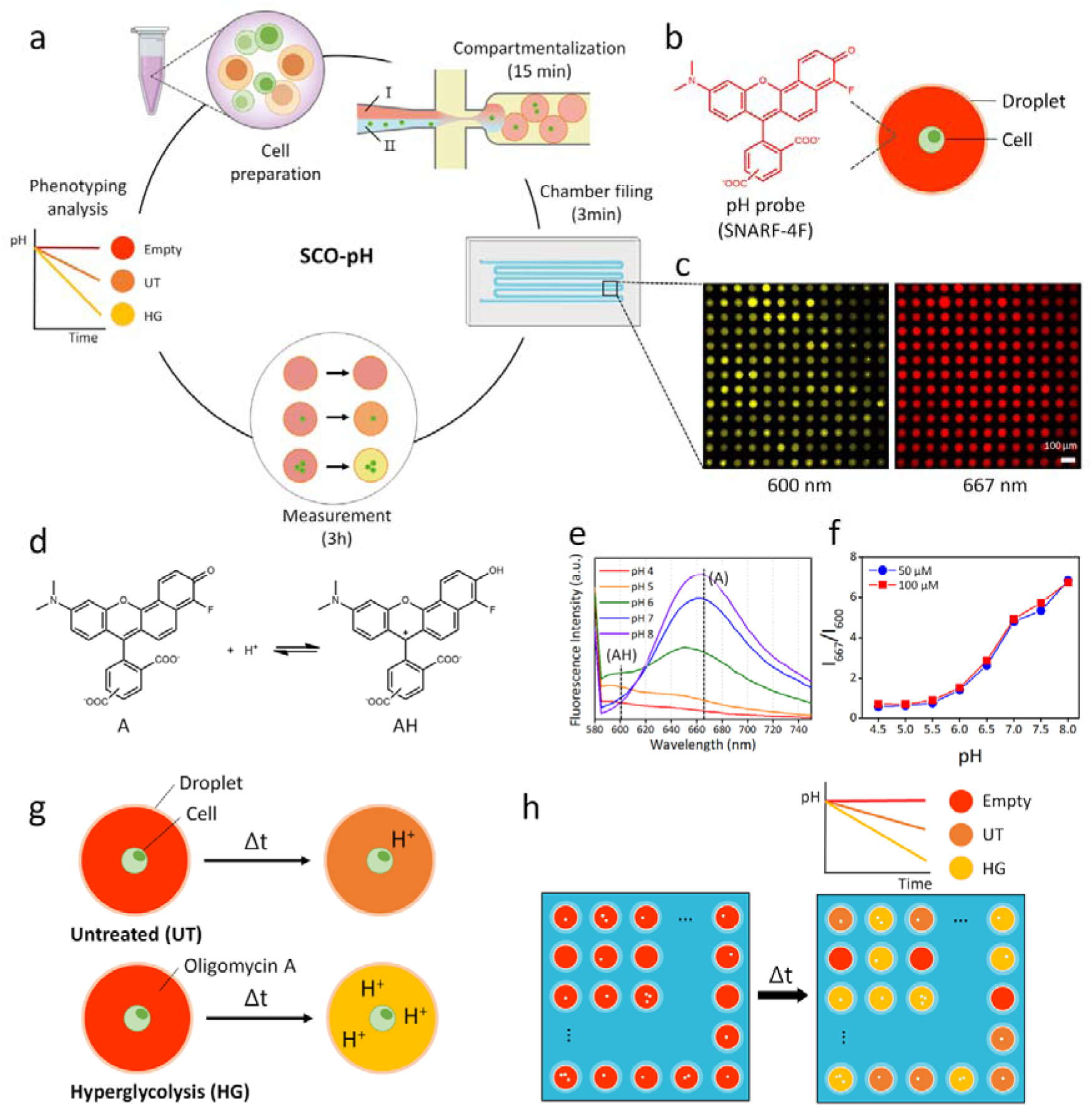
Principle and calibration of droplet-based single cell optical pH analysis platform (SCO-pH). (a) Overview of the workflow and timing: EL4 cells are cultured and compartmentalized into 143-pl droplets using a microfluidic device. Two aqueous phases (Phase □ and Phase □) are utilized, with Phase □ containing SNARF-4F (a pH probe), as a fluorescently labeled detection reagent, and Phase □ containing the cells. After generation, droplets are loaded into a microwell array for observation. Under static conditions without flow, extracellular pH and acidification rates are measured over time. A total of 40,000 droplets are observed through repeated imaging loops, displaying fluorescent intensities by incorporating SNARF-4F and cells stained with dyes. The custom MATLAB code tracks and collects intensities of both droplets and cells. (b) Droplet comprises single cell or multiple cells and a SNARF-4F. (c) Fluorescence images taken at 600 nm and 667 nm within a single imaging window, comprising 169 droplets among 240 positions in the array, show two emission intensities of SNARF-4F. (d) The acid (AH) and base (A) forms of SNARF-4F at ground-state equilibrium. (e) Fluorescence intensities of 50 μM of SNARF-4F range from pH 4 to 8. The AH form exhibits an emission peak at 600 nm, while the A form emits at 667 nm. (f) The ratio of fluorescence intensities at 600 nm (I_600_) and 667 nm (I_667_) as a function of pH indicates a correlation between I_667_/I_600_ ratio and pH, independent of SNARF-4F concentration. (g) Schematic illustrations depicting changes occurring in droplets containing untreated (UT) and hyperglycolytic (HG) single cells. (h) Representation of a microwell array with droplets displaying varying acidification levels.

To measure the extracellular pH within each droplet, we co-encapsulated cells with a cell-impermeable pH probe, SNARF-4F, in the droplets (**Fig. 1b**). SNARF-4F exists in acid (AH) and base (A) forms, with emission peaks at 600 nm and 667 nm, respectively (**Fig. 1d**). SNARF-4F has different fluorescence intensities at these two wavelengths depending on the pH condition (**Fig. 1e**), and the ratio of fluorescence intensities at 600 nm and 667 nm provides an accurate measurement of the pH of a solution (**Fig. 1f**). This ratiometric method helps control multiple artifacts in fluorescence measurements, including instrument stability and non-uniform indicator loading. To improve the accuracy of the data, we set up band-pass filters centered at 600 and 667 nm to measure the intensities of two emissions of SNARF-4F as shown in **Fig. 1c**. Additional information on the filter sets and SNARF-4F is provided in the Supplementary information (**Fig. S4**). We use a stage-top environmental chamber to maintain the cell culture environment at 95% humidity and 37°C. The configuration of the inverted microscope covered with the environmental chamber is shown in **Fig. S5a** and **S5b** in the Supplementary information.

To test extracellular acidification monitoring at the single-cell level, we developed a hyperglycolytic (HG) cell, which exhibits overactivated glycolysis upon treatment with oligomycin A in the droplets (**Fig. 1g**). The droplets containing untreated cells and those containing the HG cells are mixed and captured in the microwell array. The fluorescence intensity ratio of I_667_ and I_660_ from each droplet changes depending on extracellular pH, which is influenced by glycolysis and mitochondria-derived acidification of the two cell types (**Fig. 1h**).

### 3.2. Buffer and CO_2_ condition for extracellular pH measurement

To accurately measure the extracellular pH of each droplet, it is critical to ensure that environmental factors do not affect the medium pH and at the same time the pH probe respond sensitively to the pH change caused by cell metabolism. The pH of culture media in conventional cell culture experiments is maintained around 7.4 by the equilibrium between NaHCO_3_ in the liquid media and 5-6% CO_2_ in the gas phase.[33] We, however, find that this conventional approach is problematic because the pH of droplets devoid of cells decreases rapidly when the CO_2_ concentration is maintained at 5% in the buffered solution (**Fig. S6**). To address this issue, we leverage pH measurement capabilities and utilize a medium condition optimized for the Agilent Seahorse (Santa Clara, CA). This system employs 0% CO_2_ condition during extracellular pH measurement, along with their buffer system (Seahorse XF medium) which is free of bicarbonate and contains no phenol red. Because phenol red absorbs wavelengths of 450 and 560 nm, it overlaps with the pH probe and might affect the sensitivity of pH measurement. We maintain the CO_2_ concentration within the environmental chamber at 0%, which is continuously tracked using the monitoring system (**Fig. S5c**). According to the supplier, this buffer solution can be used for up to 6 hours and eliminates potential problems that may impact the cellular metabolism.

To determine the suitability of using this Seahorse buffer system for measuring a subtle change in the pH in these pico-scale droplets, we compared phosphate buffered saline (PBS), live cell imaging solution (LCIS), and the Seahorse XF RPMI medium for droplet-based extracellular pH measurement. The compositions of three different buffers are provided in the Supplementary information (**Table S1**). To test these three buffers, we measured fluorescence intensities of SNARF-4F in four different conditions, including 0.1X PBS, XF medium, XF medium with supplements (containing GlutaMax, glucose, and pyruvate following the pH measurement condition of Seahorse technology)[34, 35], and LCIS (0% CO_2_, 37°C, 95% humidity). We find that the fluorescence intensities remain most stable in the supplemented XF medium compared to other buffers (**Fig. S7**).

### 3.3. Droplet crosstalk and stability

To ensure that cell metabolism in a particular droplet affects only the pH of that droplet and not any others, it is crucial that the water droplets retain SNARF-4F without compromising emulsion stability and that the probe does not transfer between droplets under the experimental conditions of 37°C, 95% humidity, and 0% CO_2_. EA-surfactant is a non-cytotoxic triblock copolymer composed of poly (ethylene glycol) and perfluoropolyether (PEG-PFPE_2_) and is widely used to stabilize aqueous droplets to encapsulate cells (**Fig. 2a**). However, poor quality control during manufacturing [36, 37] can compromise the stability of EA-surfactant stabilized emulsions; moreover, small molecules (∼200 to ∼500 Dalton) can pass between EA-stabilized droplets due to the presence of micelles in the oil phase.[38–40]

**Figure 2.**
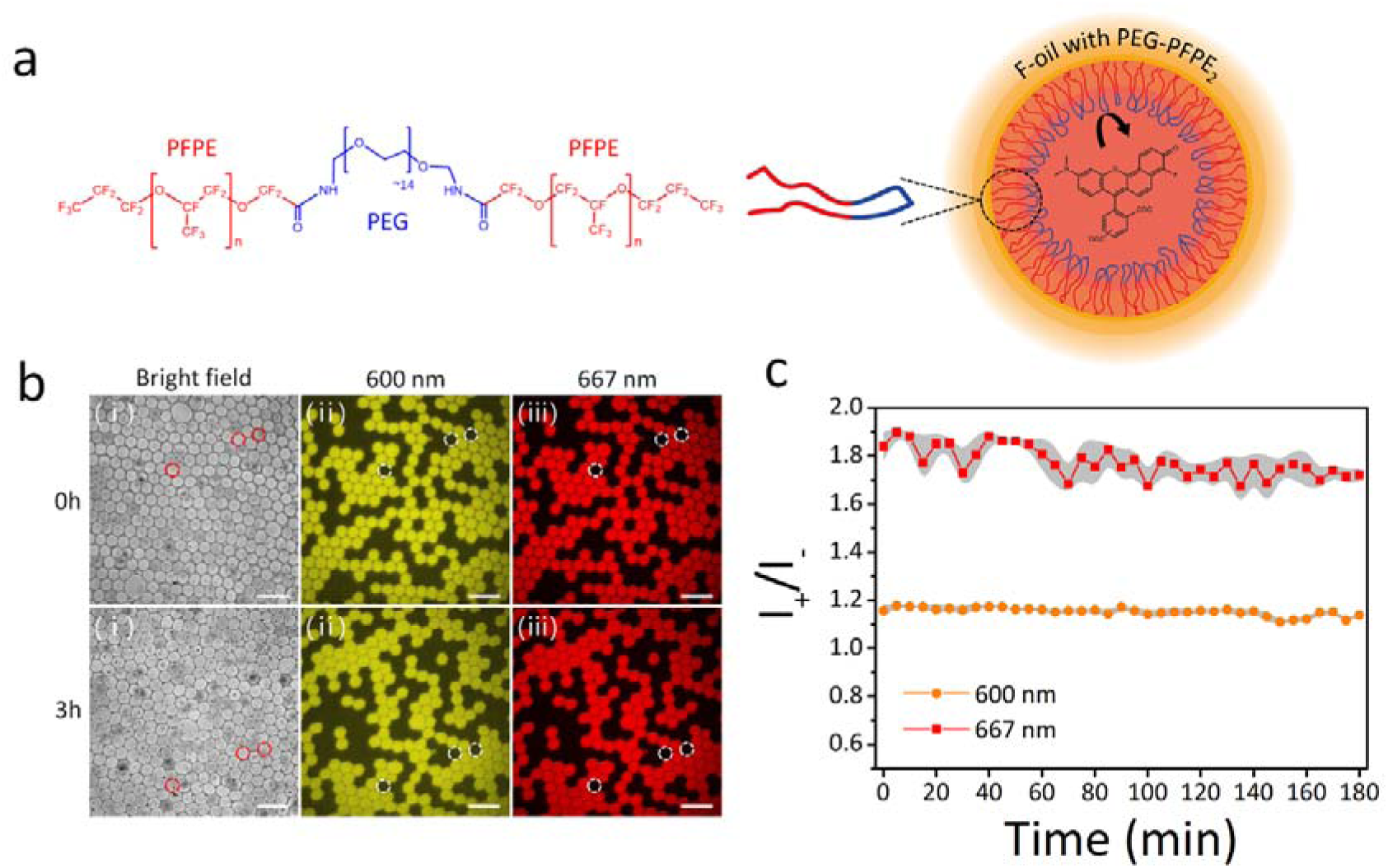
Droplet crosstalk and stability test. (a) Chemical structure of EA-surfactant, and schematic illustration of EA-stabilized droplet in the fluorinated oil phase containing SNARF-4F. (b) Fluorescence images of “positive” and “negative” droplets stabilized by EA-surfactant after mixing (0h) and after 3h. The emulsion contained a 1:1 mixture of “positive” droplets containing SNARF-4F (50 μM) and “negative” droplets containing buffer only. (□) Bright-field image, fluorescence images at (□) 600 nm and (□) 667 nm. Three negative droplets are marked with red and white dashed circles. Scale bar is 200 μm. (c) Time evolution plots of the ratio of fluorescence intensity of positive droplets (*I*_+_) versus that of negative droplets (*I*_-_) at 600 nm and 667 nm. The mean value is calculated from random three droplets in each image and the grey area represents standard deviation of the mean.

To test whether these issues could complicate our approach, we performed experiments to test the stability of emulsion droplets and the possible presence of probe transfer between droplets. We produced and mixed two types of droplets: “Positive” droplets containing SNARF-4F and “negative” droplets with buffer only. These two types of droplets are mixed a 1:1 volume ratio and observed in a PDMS-based chamber for 3 h (**Fig. 2b**). Three negative droplets are marked with a red dashed circle in the bright field image and white dashed circles in the fluorescent images at 600 nm and 667 nm. Negative droplets remain empty and stable with no size, shape, and color changes, indicating no crosstalk among the droplets stabilized with EA surfactant. The leakage of SNARF-4F is characterized by measuring the fluorescence intensity ratio (*I*_+_/*I*_-_) between SNARF-4F in positive and negative droplets over time (**Fig. 2c**). The ratio (*I*_+_/*I*_-_) indicates that two different forms of SNARF-4F remain stable in droplets stabilized by EA-surfactant in HFE-7500 for 3 h.

### 3.4. Cell model with over-activated glycolysis

To determine that SCO-pH accurately monitors the extracellular acidification of single cells, we over-activated glycolysis in cells by using oligomycin A. This is an inhibitor of ATP synthase and significantly suppresses mitochondrial respiration and maximizes glycolysis by blocking the proton channel of ATP synthase.[41] We expect that oligomycin A-treated cells would exhibit high levels of lactate production, resulting in rapid extracellular acidification of the aqueous droplets (**Fig. 3a**). This hyperglycolytic (HG) cell model is compared to the control group (UT; oligomycin A untreated group).

**Figure 3.**
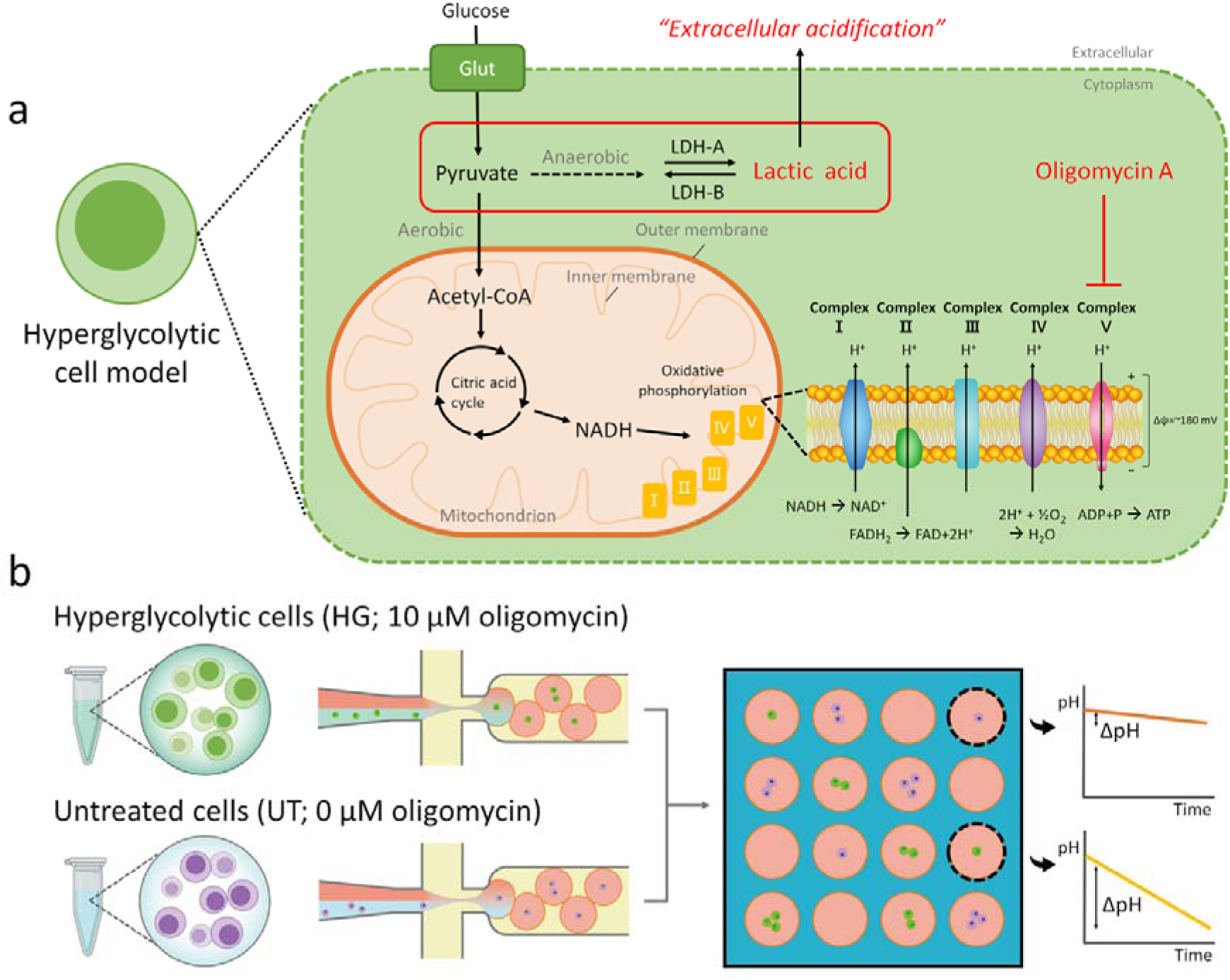
Oligomycin A-induced hyperglycolytic cell model and experimental design. (a) The hyperglycolytic (HG) cell model induced by oligomycin A, and cellular mechanism of extracellular acidification. (b) Experimental design for distinguishing HG cells from untreated (UT) cells in a microwell array. The HG cells exhibit green fluorescence, while the UT cells display violet fluorescence. Each cell type is compartmentalized separately into droplets, which are then mixed and randomly loaded into the microwell array for observation.

HG cells are prepared by staining EL4 cells with Calcein AM and co-encapsulating them with 10 μM of oligomycin A and 50 μM of SNARF-4F as shown in **Figure 3b**. To distinguish the droplets containing UT cells from those containing HG cells, EL4 cells are stained with CellTrace violet and encapsulated without oligomycin A. Two different droplets are mixed in a 1:1 volume ratio and randomly immobilized into the microwell array. With this approach, we can collect the acidification dynamics of each cell using two different sets of images, tracking only green-labeled cells or violet-labeled cells.

### 3.5. pH measurement by droplet image analysis

Images of the captured droplets in the microwell array are automatically taken by the motorized microscope stage operated by a custom script of Metamorph software. The image analysis pipeline is shown in **Fig. 4a**. Cell tracking in the droplets is performed by the free software FIJI with a Particle Tracker customized for automatic image analysis. To monitor the extracellular pH of individual droplets, we employ custom MATLAB scripts. These scripts facilitate droplet indexing and enable the discrimination between droplets containing cells and empty droplets based on cell tracking data. The results of this image analysis enable the identification of relevant droplets within the image and the tracking of their pH variations. Our MATLAB script can be tailored to generate graphical representations, including plots depicting droplet intensities, cell intensities, and extracellular pH values.

**Figure 4.**
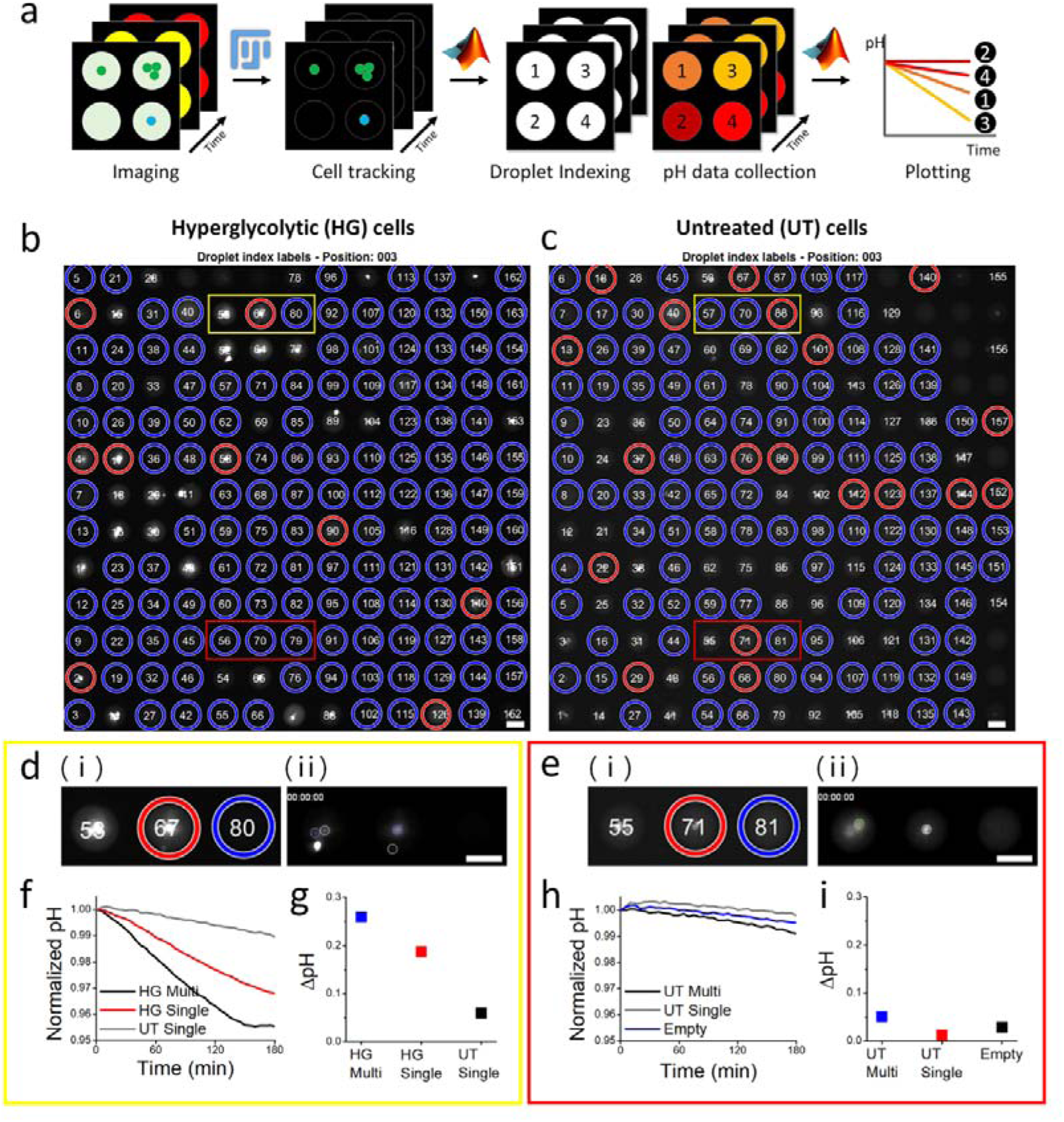
Droplet indexing and extracellular pH collection. (a) Workflow of image analysis using the ‘Particle Tracker’ of FIJI for cell tracking, followed by droplet indexing, extracellular pH collection, and plotting using custom MATLAB codes. Droplet indexing results obtained from MATLAB: (b) Hyperglycolytic (HG) cells are detected in the green channel, and (c) untreated (UT) cells are detected in the UV channel. Droplets containing a single cell are represented by red circles, empty droplets by blue circles, and droplets containing multiple cells are unmarked. For examples of pH monitoring performance, three droplets within the yellow and red boxes in both images are magnified in (d) and (e), respectively. (d) (□) Magnified images of droplet indexing for HG cells and UT cells, along with (□) images of detected cells by the Particle Tracker, are displayed. Single cells detected in the representative image at the start of imaging (0h) are outlined. (e) (□) Magnified images of droplet indexing for HG cells and UT cells are shown in red boxes. (□) The image of detected cells obtained by the Particle Tracker is displayed. Cells are outlined in the image captured at the start of imaging (0h). Color coding for (f) to (i): HG single cell droplet (red), HG/UT multiple cell droplet (black), UT single cell droplet (grey), and empty droplet (blue). (f) Normalized pH and (g) raw ΔpH (initial pH-final pH; pH_i_-pH_f_) for the droplets shown in (d) over 3 hours. (h) Normalized pH and (i) raw ΔpH (initial pH-final pH; pH_i_-pH_f_) for the droplets shown in (e) over 3 hours. The scale bar is 50 μm.

The droplet indexing protocol identifies droplets containing single cells with red outlines and empty droplets with blue outlines. Droplets with no outlines represent multiple cell droplets. Droplet indexing outputs in a representative position confirm that the image pipeline reliably works for both images regardless of which dye the cells are stained with (**Fig. 4b** and **4c**). As an example, we observe three droplets in a yellow box that has been placed in an arbitrary location for both HG and UT cells (**Fig. 4d**). The image of outlined cells by the Particle Tracker (**Fig. 4d**, (□)) is displayed next to the droplet indexing image (**Fig. 4d**, (□)). In the droplet containing single cell (middle), the droplet boundary is detected as a cell. We find that the Particle Tracker is prone to error near the outer boundaries of each image as it misidentifies high contrast patches at the edge of droplets as cells. Therefore, we analyze the central 60% of the droplet area to identify the number of cells in each droplet. Upon applying the code to the 6 droplets in the yellow boxes (3 HG and 3 UT droplets), we successfully identify one HG multiple cell, one HG single cell, and one UT single cell (**Fig. 4d, Fig. S8**). As shown here, we can find out that empty HG droplet is not actually empty and is a UT single cell droplet by comparing two images. In another location marked using red boxes, no HG cells are found, and only UT cells are detected. The first droplet contains UT multiple cells, and the second droplet contains a UT single cell. The third droplet is empty as it appears empty in both images (**Fig. 4e**). We provide cell tracking videos of those droplets in the Supplementary information (**Video S1**).

To verify the validity of SCO-pH for extracellular pH monitoring of single cells, we characterize the pH curves from each droplet based on the MATLAB outputs. The first location (yellow box) shows different acidification dynamics among the droplets. A HG single cell droplet shows drastic extracellular pH reduction over time compared to a UT single cell droplet. Also, the HG multiple cell droplet shows faster acidification than the HG single cell droplets, as expected (**Fig. 4f**). The actual pH of the droplets can be calculated based on a calibration curve created by an equation provided by the manufacturer of SNARF-4F (See supplementary information in **Fig. S9**). To verify how much acidification occurs in the droplet by cells, we compare raw ΔpH (initial pH-final pH) of each droplet based on the pH calculation (**Fig. 4g**). Raw ΔpH of each droplet containing the HG multiple cells and the HG single cell decreases by 0.26 and 0.19, respectively. In contrast, the UT single cell droplet shows only 0.06 raw pH shift in 3 hours. Another example in a different location (red box) clearly shows that in the case of the UT cell droplets, extracellular acidification is not different between the multiple cell droplet and the single cell droplet. Normalized pH plots show that all plots have similar trends among the UT single cell droplet, the UT multiple cell droplet, and the empty droplets (**Fig. 4h**). Extracellular raw pH decreases by only 0.05, 0.01, and 0.03, respectively (**Fig. 4i**).

### 3.6. Correlation analysis between extracellular pH and glycolysis

Our results above demonstrate that SCO-pH detects extracellular pH in droplets reliably at single cell levels. Additionally, our custom image analysis pipeline rapidly and accurately tracks the pH of droplets and classifies droplets. Using SCO-pH, we analyze all the droplets in three positions out of 240 positions in the entire microwell array to validate the acidification monitoring performance at the single cell level. A total of 507 droplets are analyzed from the three positions. In these areas, single cell encapsulation rate is 17.6%, less than calculation (30%). We observed two types of cells with green and violet fluorescence, and some highly fluorescent cells generate outlier errors in the data. After eliminating errors, the single cell encapsulation rate that can be identified through our code is 11.8%.

We compare the extracellular acidification of single and multiple cells in the HG and the UT cell models. The normalized pH plots of droplets containing HG cells show different acidification patterns among single cell droplets, multiple cell droplets, and empty droplets (**Fig. 5a**). Also, the pH curve distribution of individual droplets indicates compartmentalized patterns (**Fig. 5b**). Multiple cells decrease the pH of the droplet more rapidly compared to single cells. In comparison of ΔpH, the extracellular pH of single cell droplets decreases by 0.17±0.07 units over a span of 3 hours. This value is 8 times higher than that for empty droplet and 0.7 times lower than that for multiple cell droplets. ΔpH of single cell droplets is also significantly different from multiple cell droplets and empty droplets (**Fig. 5c**).

**Figure 5.**
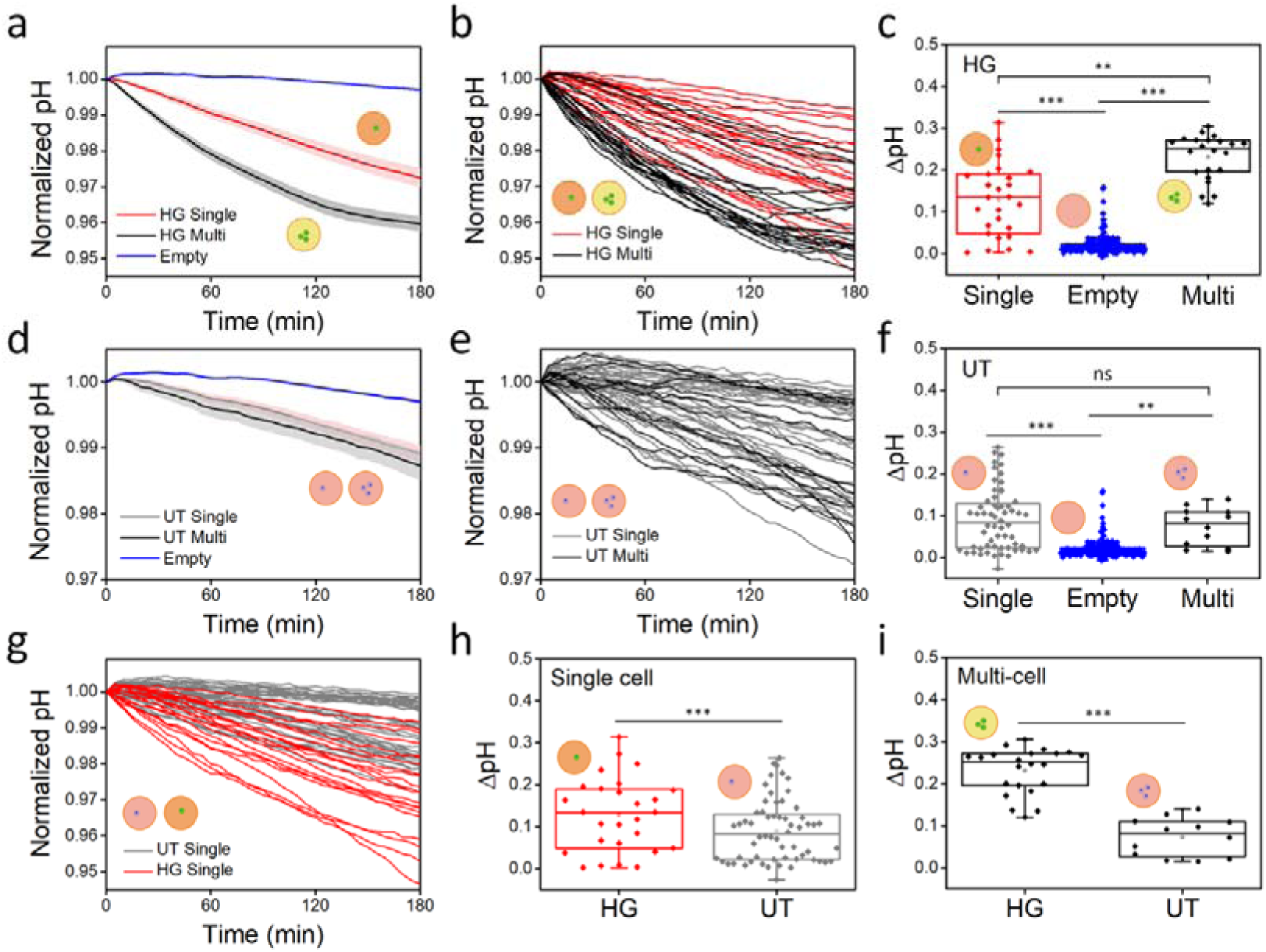
Extracellular pH differences in cell models with different acidification observed for 3 hours in three different positions in a device. The hyperglycolytic (HG) cell model analysis: (a) Normalized and averaged extracellular pH of the droplets containing single cells (red), multi cells (black), and no cells (empty; blue) tracked for 3 hours. Shaded regions correspond to standard error. (b) Individual normalized extracellular pH curves of the droplets with single cells and multi cells. (c) Raw ΔpH (initial pH-final pH; pH_i_-pH_f_) of the droplets containing single cells, multi cells, and no cells for 3 hours. The untreated (UT) cell analysis: (d) Normalized and averaged extracellular pH of the droplets containing single cells, multi cells, and no cells tracked for 3 hours. Shaded regions correspond to standard error. (e) Individual normalized extracellular pH curves of the droplets containing single cells and multi cells. (f) Raw ΔpH of the droplets containing single cells, multi cells, and no cells for 3 hours. Comparison of the HG single cell droplets (red) and the UT single cell droplets (gray): (g) Normalized extracellular pH curves of the individual droplets containing the HG and the UT single cells. (h) Raw ΔpH of the droplets containing the HG and the UT single cells. (i) Raw ΔpH of the droplets containing the HG and the UT multiple cells. The significance of the data is calculated by the paired Student’s *t*-test (**, p <0.01; ***, p < 0.001).

In the case of the UT cells, single cell droplets and multiple cell droplets are not significantly different in the averaged pH curves and individual pH curve distribution (**Fig. 5d** and **5e**). Nonetheless, single cell droplets and multiple cell droplets differ significantly from empty droplets due to glycolysis (**Fig. 5f**). Given the absence of disparity in acidification between single cells and multiple cells, the impact of acidification on UT cells appears to be minimal.

Finally, we compare the ΔpH of the droplets contain single cells to assess the utility of this technology in analyzing the heterogeneity of individual cells. The pH distribution of single cell droplets reveals different patterns for HG and the UT cells (**Fig. 5g**). Generally, the red plots representing HG single cell droplets exhibit a more rapid decreasing trend compared to the grey plots of UT single cell droplets. Nevertheless, individual cells have different acidification dynamics with fair variation in the rates of acidification, including some HG cells showing slower decrease than control cells, representing single cell heterogeneity in metabolism. These results indicate the capability of SCO-pH to measure extracellular acidification at single cell level and assess population heterogeniety. Overall, extracellular pH change of HG single cell is 0.17 in 3 hours, which is 2.6 times higher than the one for the UT cells (**Fig. 5h**). In the case of multiple cell droplets, the extracellular pH change of the HG is 0.23 in 3 hours, 3.2 times higher than that of the UT cells (**Fig. 5i**).

### 3.7. Extracellular pH monitoring performance of the entire device

Finally, we analyze the extracellular pH dynamics of EL4 cells in the glucose-rich environment (10 mM) across all positions in the microwell array (239 positions). To quantify the error rates of this technology, we manually find errors (i.e., those identified as “single cells” containing multiple cells or no cell, or those identified as “empty” containing one or more cells) in Position 1 and 100. The error rates in Position 1 are 15.4% and 16.0% for single cell droplets and empty droplets, respectively. In the case of single cell droplets, high values of raw ΔpH among outliers over 0.83 are identified as multiple cell droplets, and low values under 0.28 are empty droplets containing debris. For empty droplets, low values among outliers are identified as debris that is not a cell and droplets stuck in the wrong area. High values are from single cell droplets misidentified due to cells having low fluorescence intensity (**Fig. 6a**). We analyze Position 100 as another representative example to quantify the error rates in the same way. The error rates are 18.9% and 13.5% for single cell droplets and empty droplets, respectively. We find that if the ΔpH of empty droplets are greater than 0.41, it is due to single cell droplets and multiple-cell droplets containing cells with weak fluorescence intensities. For single cell droplets, low ΔpH under 0.37 is identified as debris, not cells (**Fig. 6b**).

**Figure 6.**
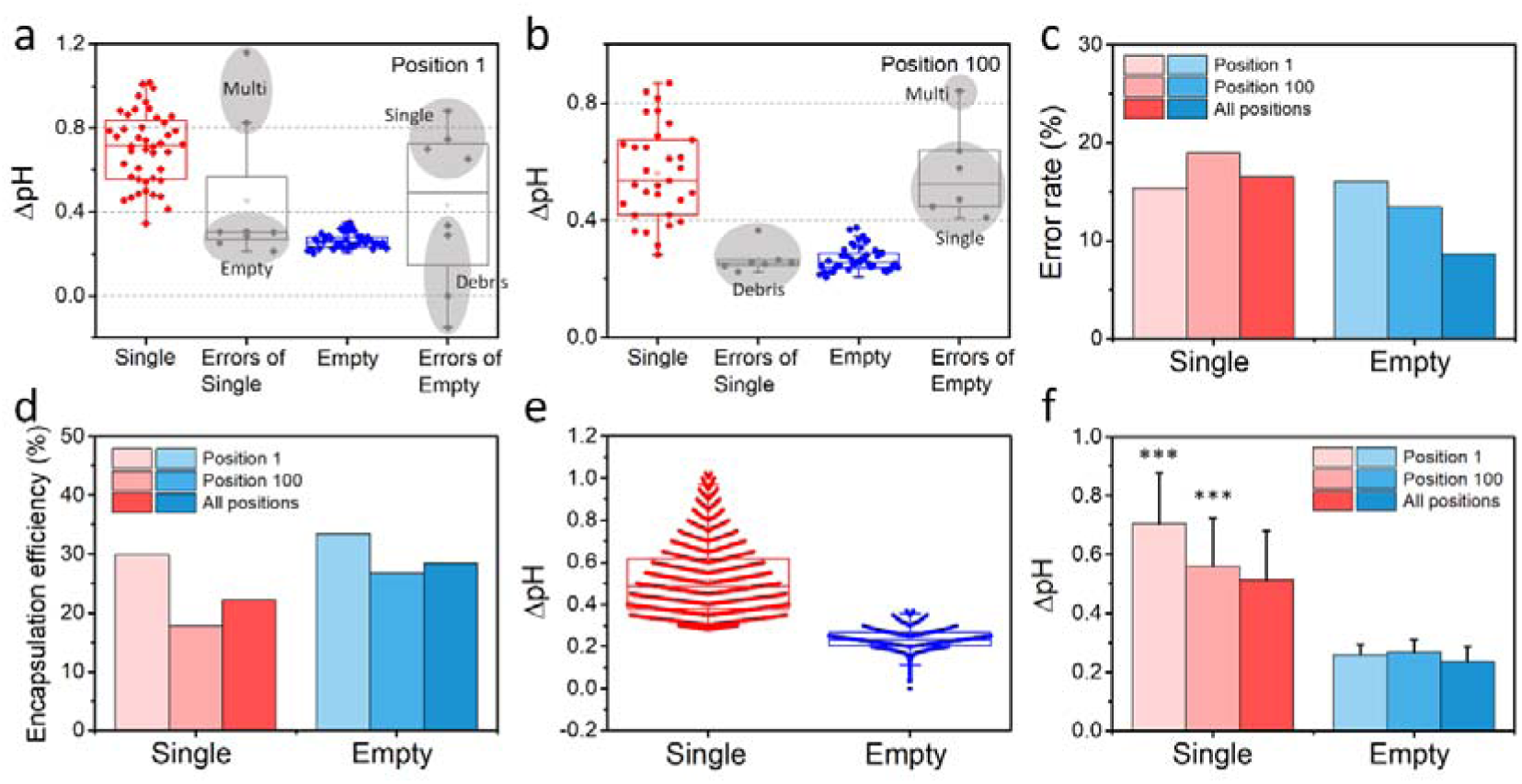
Extracellular pH monitoring performance of an entire device (239 positions): error rate, encapsulation efficiency, and extracellular pH change of single cells for 3 hours of glucose consumption. (a) Representative raw ΔpH and error distribution of single cell droplets and empty droplets in Position 1. (b) Representative raw ΔpH and error distribution of single cell droplets and empty droplets in Position 100. (c) The error rates for single cell droplets and empty droplets in Position 1, 100, and all 239 positions. (d) Encapsulation efficiency of single cell droplets and empty droplets in Position 1, 100, and all 239 positions. (e) Raw ΔpH of single cell droplets (n = 8,600 of 38,759 droplets analyzed) and empty droplets (n = 11,012 of 38,759 droplets analyzed) across all positions. (f) Raw ΔpH of single cell droplets and empty droplets in Position 1, 100, and all 239 positions. The significance of the data is calculated by the paired Student’s *t*-test (***, p < 0.001).

According to the manually analyzed results, we regard the data in the range lower than 0.37 as empty droplets and the data in the range of 0.28-1.02 as single cell droplets. After applying this criterion, we obtain 11,012 empty droplets and 8,600 single cell droplets out of 38,759 droplets in the microwell array. The error and encapsulation rates of single cell droplets are 16.5% and 22.2%, respectively. It is estimated that the debris of empty droplets causes 88.6% of the errors (ΔpH < 0.28). Only 11.4% of errors are estimated to be misidentified multi-cell droplets (ΔpH > 1.02). The error and generation rates of the empty droplet are 8.7% and 28.4%, respectively. We estimate that most errors are droplets containing cells with low fluorescence intensity (ΔpH > 0.37). The error and encapsulation/generation rates are similar to the values for Position 1 and 100 (**Fig. 6c** and **6d**). For the single cell droplet, encapsulation efficiency reaches 22.2% after eliminating the errors, and we still obtain data for over 8,000 single cells. Although the error rate is 16.5%, the remaining data is close to what we aimed for based on the calculation (30%, 12,000 single cells) and will be sufficient to determine the metabolic characteristics of single cells.

Raw ΔpH distributions of single cell droplets and empty droplets in all positions are displayed, and the two data exhibit different median values and distribution patterns (**Fig. 6e**). Furthermore, we can see the metabolic heterogeneity of single cells through a large distribution area. The averaged raw ΔpH values of single cell droplets and empty droplets of all positions are similar to the values of Position 1 and 100 (**Fig. 6f**). Raw ΔpH of single cell droplets and empty droplets are significantly different due to the acidification of cells. Taken together, we demonstrate that SCO-pH is able to perform various tasks from imaging to analysis successfully throughout the entire microwell array to track extracellular pH over time, and thus be useful for applications that require dynamic monitoring of acidification at the single cell level.

## 4. Conclusions

As a single cell phenotyping tool, SCO-pH successfully measured the temporal acidification dynamics of single cells over 3 hours from thousands of droplets. Cells are compartmentalized into pico-liter scale droplets which are captured in the two-dimensional array to provide temporal readouts. Measuring the extracellular pH in these droplets using a cell-impermeant ratiometric pH indicator can provide new insights into single cell phenotyping based on their metabolism. The pH monitoring data of thousands of droplets are sorted by the number of cells in droplets, and the data from single cell droplets show broadly different acidification patterns of HG cells and UT cells, while also revealing their single cell heterogeneity. We believe SCO-pH will be potentially useful for the detection of rare cells with abnormal glycolysis such as cancer cells and activated immune cells for disease diagnosis and patient’s physical condition monitoring.

Our system allows identification of droplets with cells exhibiting desired or unusual metabolic characteristics based on temporal dynamics. It could be extended to physically collect selected droplets that exhibit unique trends using a previously developed laser-based method[29]. Furthermore, this tool is easily scalable to observe a large number (potentially beyond 40,000) of cells at a time and can be adapted to screen other types of cells beyond the immune cells.

## Acknowledgements

Research reported in this publication was supported by the National Human Genome Research Institute of the National Institutes of Health under Award Number RM1HG010023. The content is solely the responsibility of the authors and does not necessarily represent the official views of the National Institutes of Health. E.A.L.P. is funded through the University of Pennsylvania Fontaine Society.

## Conflicts of Interest

The authors declare no conflict of interest.

## Author Contributions

H.J., S.H.H., D.L., and J.K. designed the study. H.J. implemented experiments, analyzed the data, and wrote the manuscript. E.A.L.P. and B.L. created the image analysis pipeline. D.K. wrote the image file sorting script. The other authors assisted with the experiments and discussed the results. All authors have read and approved the final version of the manuscript.

## Supporting Information

## Materials and methods

### Materials

Tetraethyl orthosilicate (TEOS) was purchased from Sigma-Aldrich (MO, USA). Ethyl alcohol 200 proof (EtOH, 99%, anhydrous) was obtained from Decon laboratories, Inc. (PA, USA). Ammonium hydroxide (NH_4_OH, 28-30%) was obtained from Fisher Scientific (NH, USA). 1H,1H,2H,2H-Perfluorooctyltriethoxysilane (FAS) was purchased from Alfa Aesar (MA, USA).

### Synthesis of Silica Nanoparticles

70 nm of silica nanoparticles (SiO_2_ NPs) were prepared based on the modified Stöber synthesis method reported in the literature. [1–3] TEOS, EtOH, and NH_4_OH were used as materials. 1.5 mL of TEOS was quickly added to 50 mL of 6%(v/v) NH_4_OH solution in EtOH with vigorous stirring. The solution was stirred for 12 h at the ambient condition (25°C, 1 atm). Synthesized SiO_2_ NPs were washed three times with EtOH using centrifugation (11,000 rpm, 15 min) and the pellet was dispersed in EtOH without drying.

### Fluorination of Silica Nanoparticles

Fluorination of SiO_2_ NPs was conducted by following the previously reported method. [4] As-prepared SiO_2_ NPs were dispersed in EtOH at 5.75 mg/mL and 153.6 μL of NH_4_OH was added to 5.37 mL of SiO_2_ NPs solution. Afterwards, 948 μL of FAS was quickly injected into the solution with vigorous stirring. The volume of FAS was calculated by comparing the specific surface area (S_0_) of the synthesized SiO_2_ NPs and the reported particles based on equation (1).

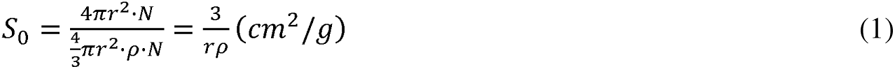

Where r, p, and N are the radius, density, and the number of SiO_2_ NPs in 1 g, respectively.

After 40 min reaction at the ambient condition (25°C, 1 atm), the particles were washed three times with EtOH using centrifugation (11,000 rpm, 15 min) and the pellet was dried overnight in the oven at 80°C. Fluorinated SiO_2_ NPs were highly dispersible in fluorinated solvents and prepared at 2%(w/w) in HFE-7500. It was mixed with FluoSurf, which is HFE-7500 containing 2%(w/w) EA surfactant (PEG-PFPE amphiphilic block copolymer, RainDance Technologies) [5], in a 1:1 ratio and used as a surfactant for Pickering emulsion.

### Characterization of Silica Nanoparticles

Morphologies of the SiO_2_ NPs before and after fluorination were observed by environmental scanning electron microscopy (SEM; FEI Quanta 600 FEG Mark II, FEI Company, OR, USA). Size and ζ-potential analysis were conducted via dynamic light scattering (DLS) (Beckman Coulter Delsa Nano C; CA, USA)(**Figure S1**).

**Figure S1.**
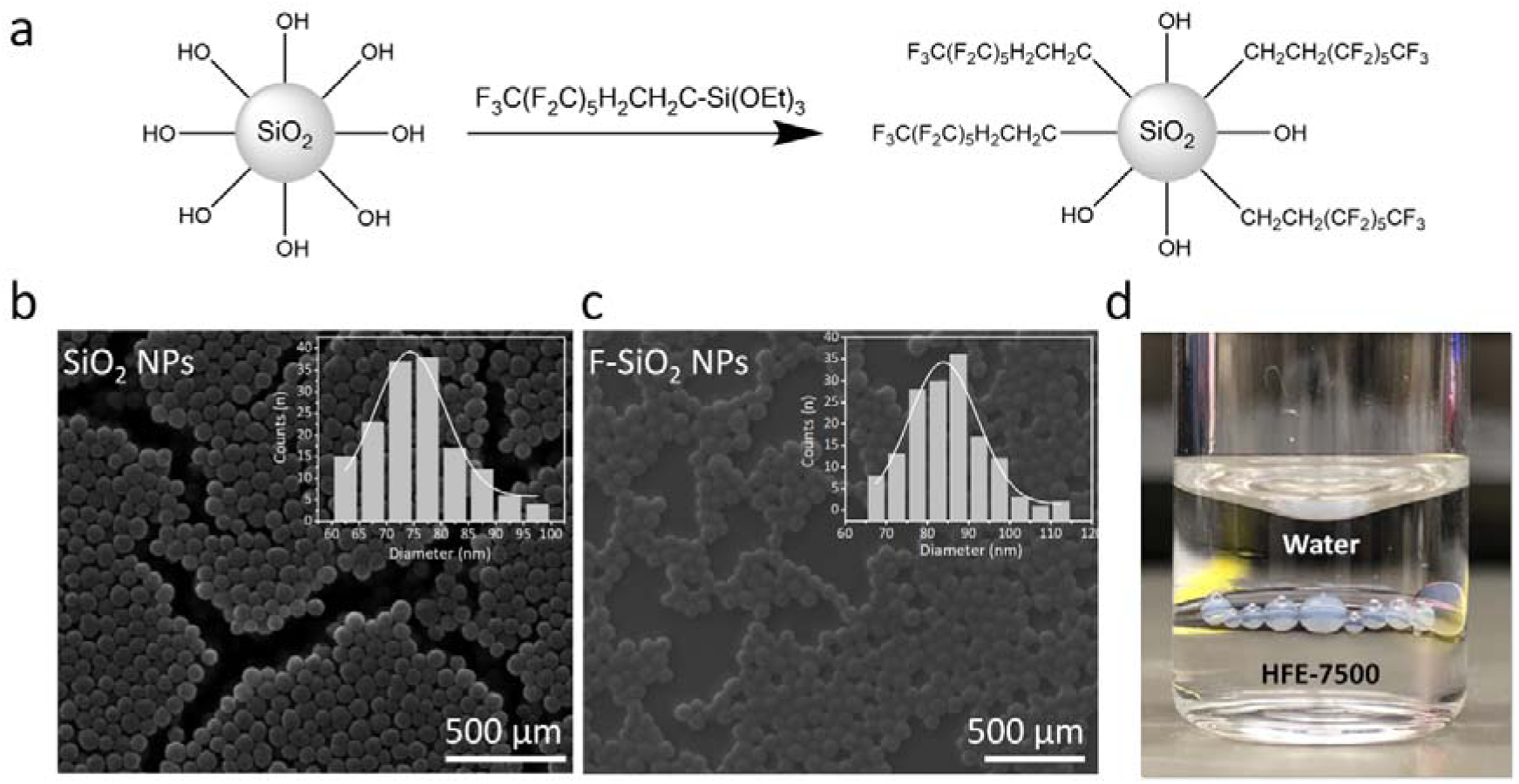
Fluorinated silica nanoparticles (F-SiO_2_ NPs) surfactant synthesis and characterization. (a) Chemistry of F-SiO_2_ NPs surfactant synthesis based on SiO_2_ NPs and 1H,1H,2H,2H-perfluorooctyltriethoxysilane (FAS), (b) Scanning electron microscope (SEM) image of synthesized SiO_2_ NPs and size distribution graph (75.6±8.4 nm, 11.2% polydispersity), (c) SEM image of synthesized F-SiO_2_ NPs and size distribution graph (83.9±9.9 nm, 11.8% polydispersity), (d) HFE-7500 containing 1 w/w% F-SiO_2_ NPs and 1 w/w% EA performs as a surfactant by locating at the interphase between water and HFE-7500.

**Figure S2.**
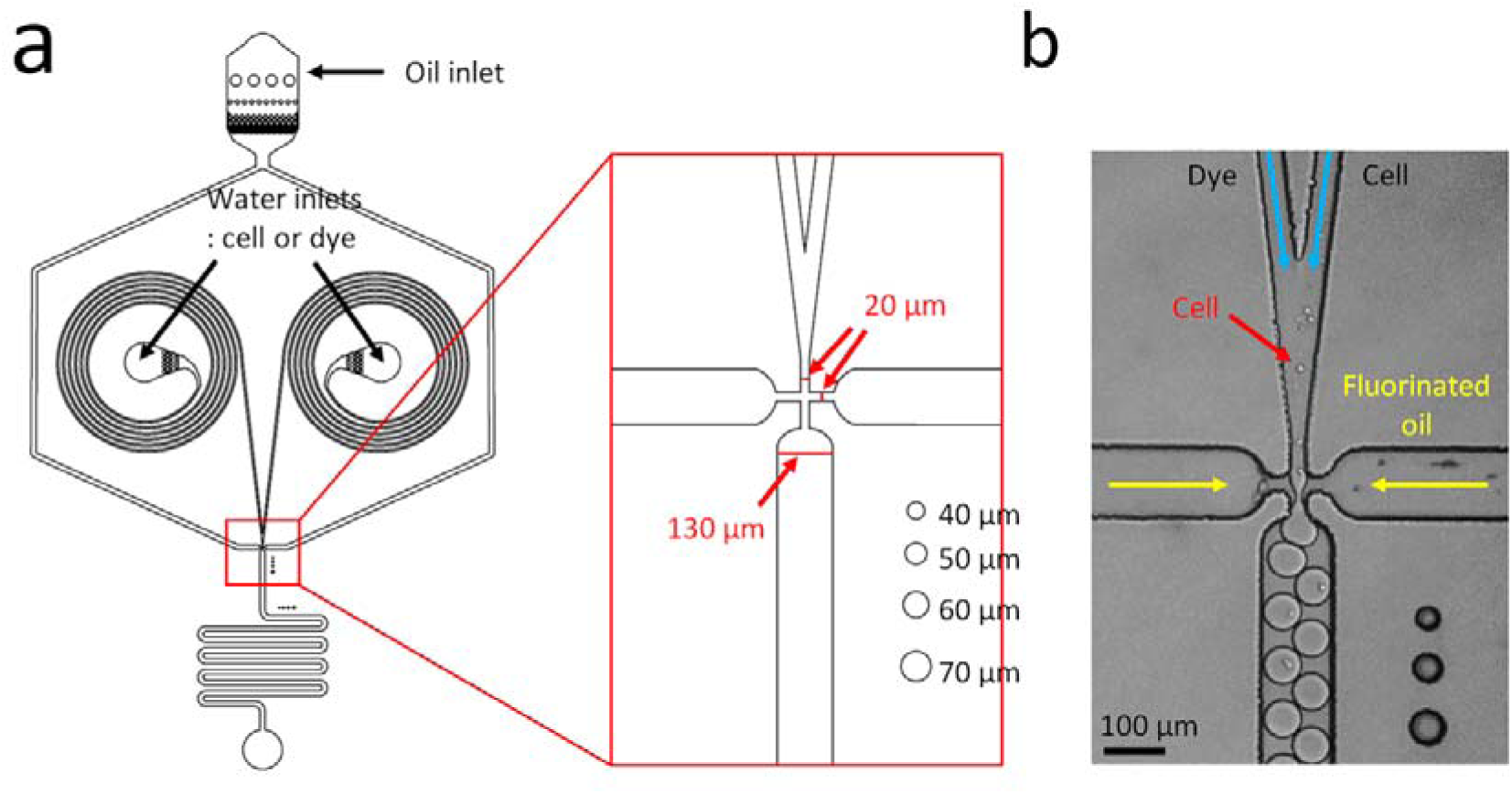
Microfluidics devices information for single cell encapsulation. (a) Droplet generator photomask design generated by AutoCAD program. The red box shows an enlarged T-junction, and the detailed channel size and scale bars for droplets are indicated in the box. (b) The image of polydimethylsiloxane (PDMS)-based droplet generator captured by a high-speed camera while droplet generation. The direction of water and fluorinated oil solutions are indicated in the image. The generator has two water inlets, one for dye and one for cells. Fluorinated oil is injected into the oil inlet with some filter features on the top of the generator. Oil comes to the junction horizontally, cutting the aqueous solution to generate droplets. The droplet generator height is 40 μm.

**Figure S3.**
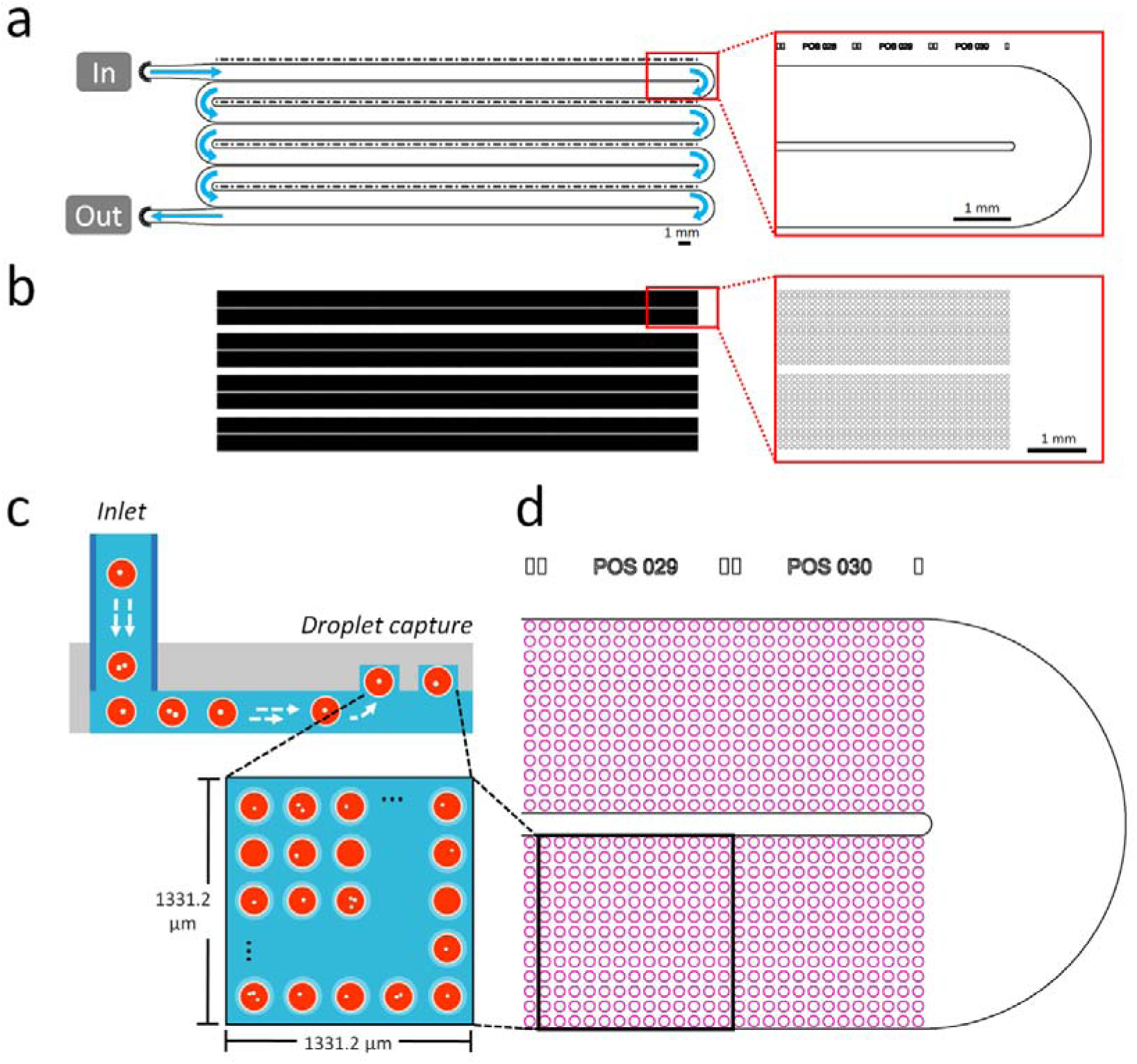
Microfluidics devices information for microwell array and imaging. Microwell array photomask design with two layers: (a) a bottom droplet flowing channel and (b) a top microwell array. The height of the droplet flowing channel is 60 μm. Inlet and outlet are displayed in the left image, and the light blue arrows indicate direction of flow. A small red box is enlarged on the right side to present two lane pair and position numbering above the first lane. (b) The height of a microwell is 50 μm (110 μm from the bottom). A small red box is enlarged on the right side to present two lane pair, and the lane containing small circles, which are microwells. (c) Schematic illustration of the cross-section of the microwell array. Droplets enter the inlet and are automatically captured into microwells above the flowing channel due to the density difference between water and fluorinated oil. The imaging window indicated by the black square contains 169 microwells (13 x 13) and is viewed from the top of the microwell. This imaging window corresponds to (d) “Position 32” on the photomask design. A small microwell array includes 240 positions for imaging.

**Figure S4.**
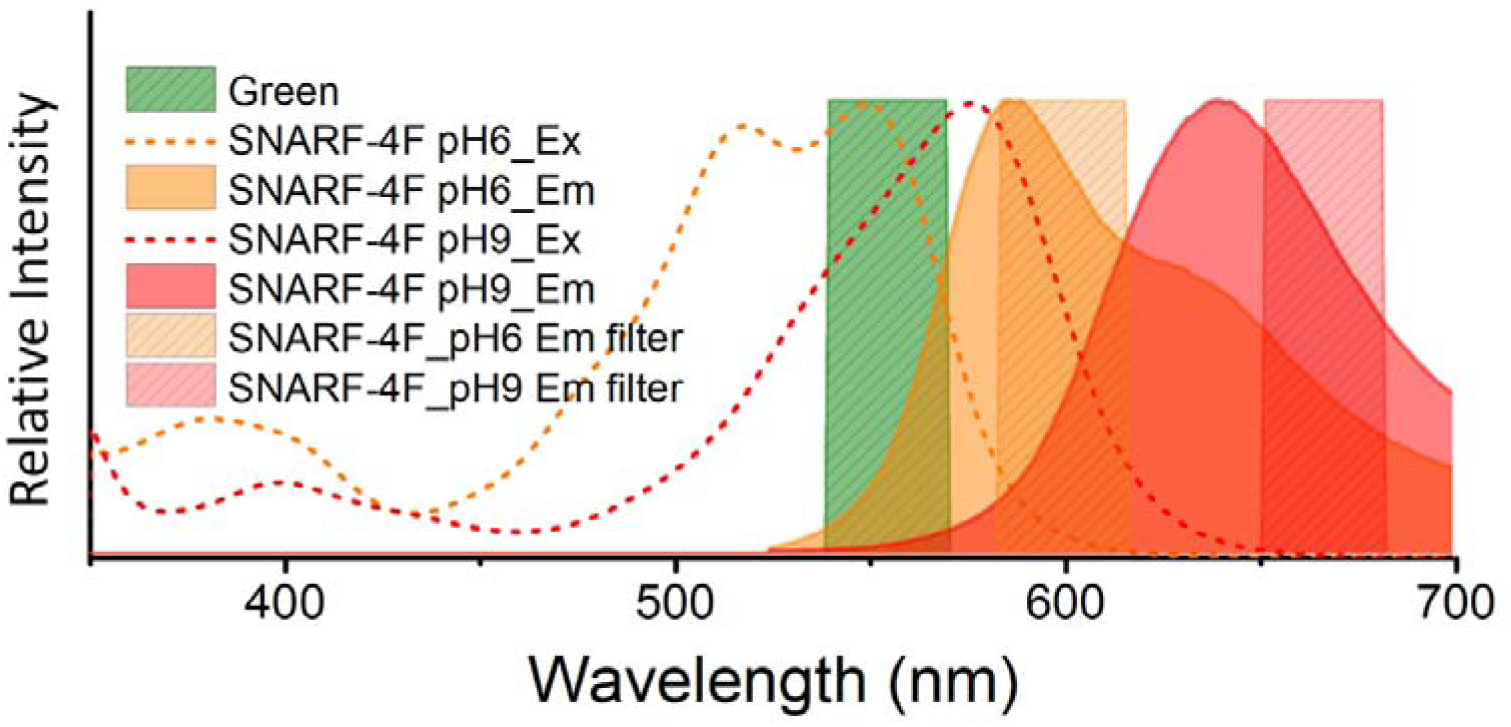
Optical information of SNARF-4F (a fluorescent pH probe), laser, and filters. Green hatched graph indicates green laser. The dotted lines are excitation plots of SNARF-4F at pH 6 (orange) and pH 9 (red). The solid lines with color-filled area are emission plots of SNARF-4F at pH 6 (orange) and pH 9 (red). The hatched graphs indicate emission filters for SNARF-4F at pH 6 (orange) and pH 9 (red). Ex, excitation; Em, emission.

**Figure S5.**
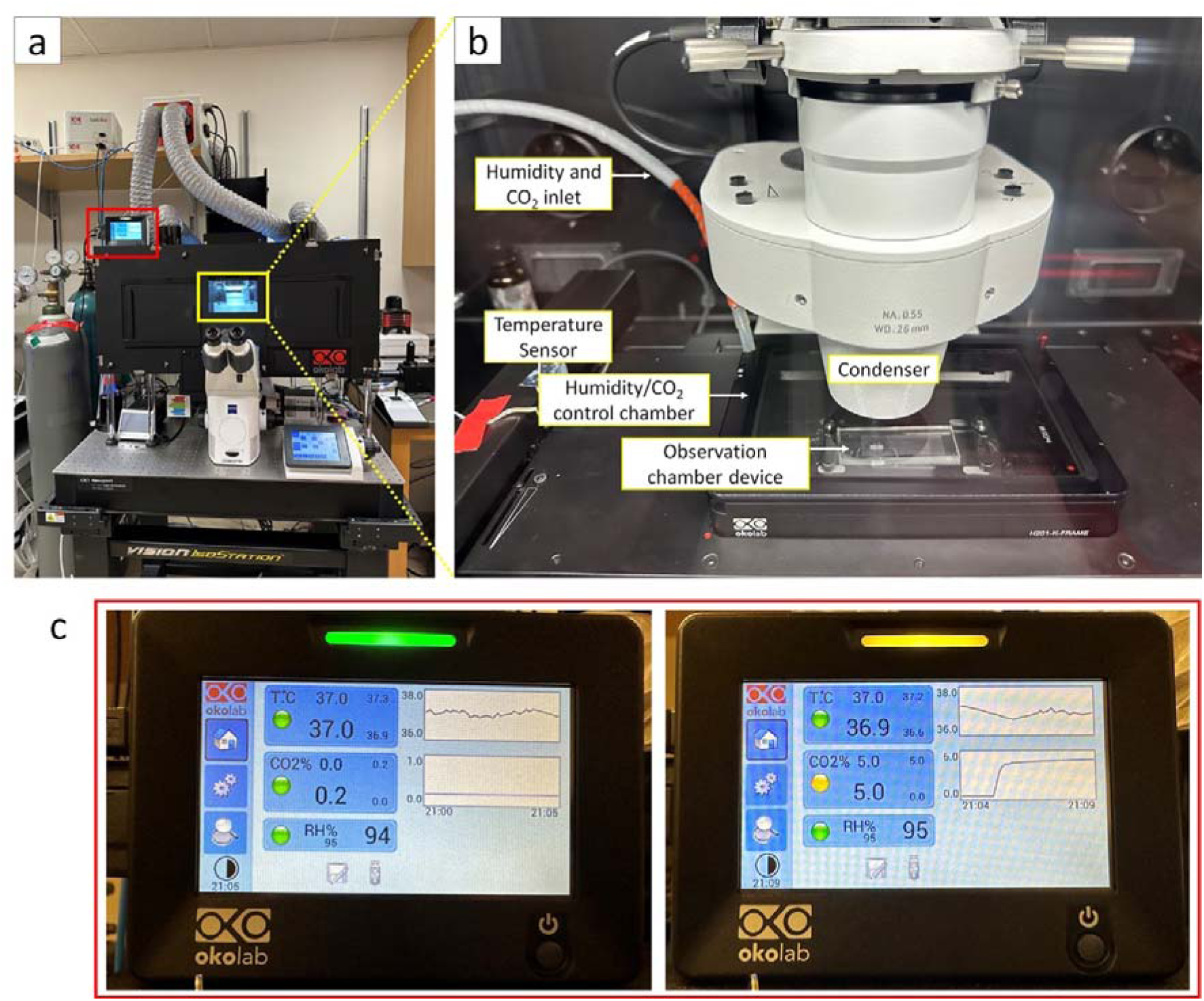
Microscope organization for single cell optical pH analysis technology. (a) The entire inverted microscope body is enclosed within an environmental chamber capable of controlling temperature, humidity, and CO_2_ concentration. (b) Inside view of the environmental chamber through the front window. The chamber maintains a temperature of 37°C and is kept dark. The temperature is monitored by a temperature sensor within the environmental chamber, while humidity and CO_2_ concentration are controlled in a separate smaller chamber located on the stage. Inlets for humidity and CO_2_ are connected to this chamber. The PDMS-based microwell array is positioned inside the humidity/CO_2_ chamber and secured to the stage using magnets. (c) The observation environment is displayed in real-time on the monitor of the environmental chamber. Changes in CO_2_ concentration from 0% to 5% can be rapidly applied to this equipment.

**Figure S6.**
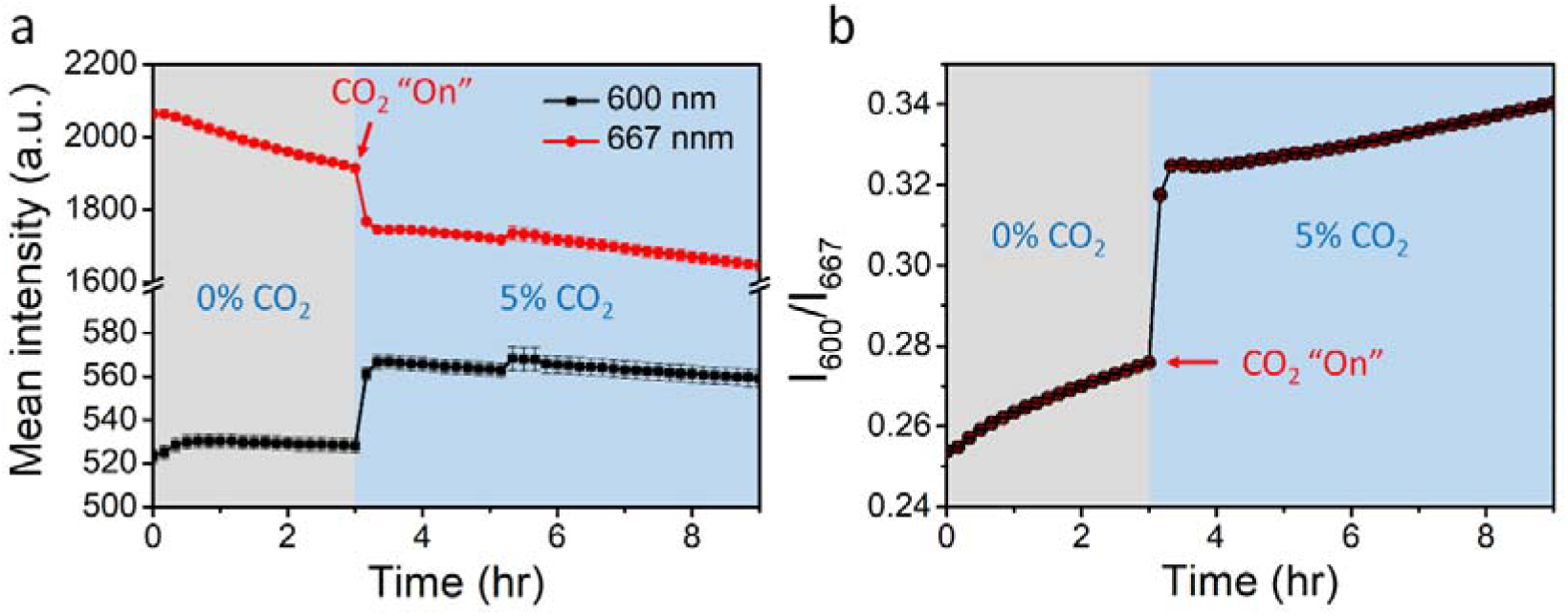
pH-reducing effect of CO_2_ on live cell imaging solution (LCIS). Droplets containing 50 μM SNARF-4F in LCIS were generated and loaded into the microwell array. The array was observed at 37°C, 95% humidity, and 0% CO_2_ for 3 hours, and then subsequently observed at 5% of CO_2_ for another 6 hours. One position out of 240 positions in the chamber was measured at 10-min intervals for a total of 9 hours. Intensities at 600 nm and 667 nm of each droplet were collected using a custom MATLAB code. (a) Mean intensity of 169 droplets at one position at 600 nm and 667 nm with standard error of the mean. (b) Mean intensity ratio between 600 nm and 667 nm for each droplet with standard error of the mean.

**Table S1.** Comparison of buffer compositions: PBS, live cell imaging solution (LCIS), and Seahorse XF RPMI medium.

| Concentration (mM) | PBS 1X (pH 7.4) | LCIS (pH 7.4) | Seahorse XF RPMI medium pH 7.4 |
| --- | --- | --- | --- |
| NaCl | 58 | 140 | 136.9 |
| KCl | - | 2.5 | 5.4 |
| HEPES | - | 20 | 1.0 |
| KH<sub>2</sub>PO<sub>4</sub> | 136 | - | - |
| Na<sub>2</sub>HPO<sub>4</sub>·7H<sub>2</sub>O | 268 | - | - |
| NaHPO<sub>4</sub> | - | - | 5.6 |
| CaCl<sub>2</sub> | - | 1.8 | - |
| MgCl<sub>2</sub> | - | 1.0 | - |
| Ca(NO<sub>3</sub>)<sub>2</sub> | - | - | 0.4 |
| MgSO<sub>4</sub> | - | - | 0.4 |

**Figure S7.**
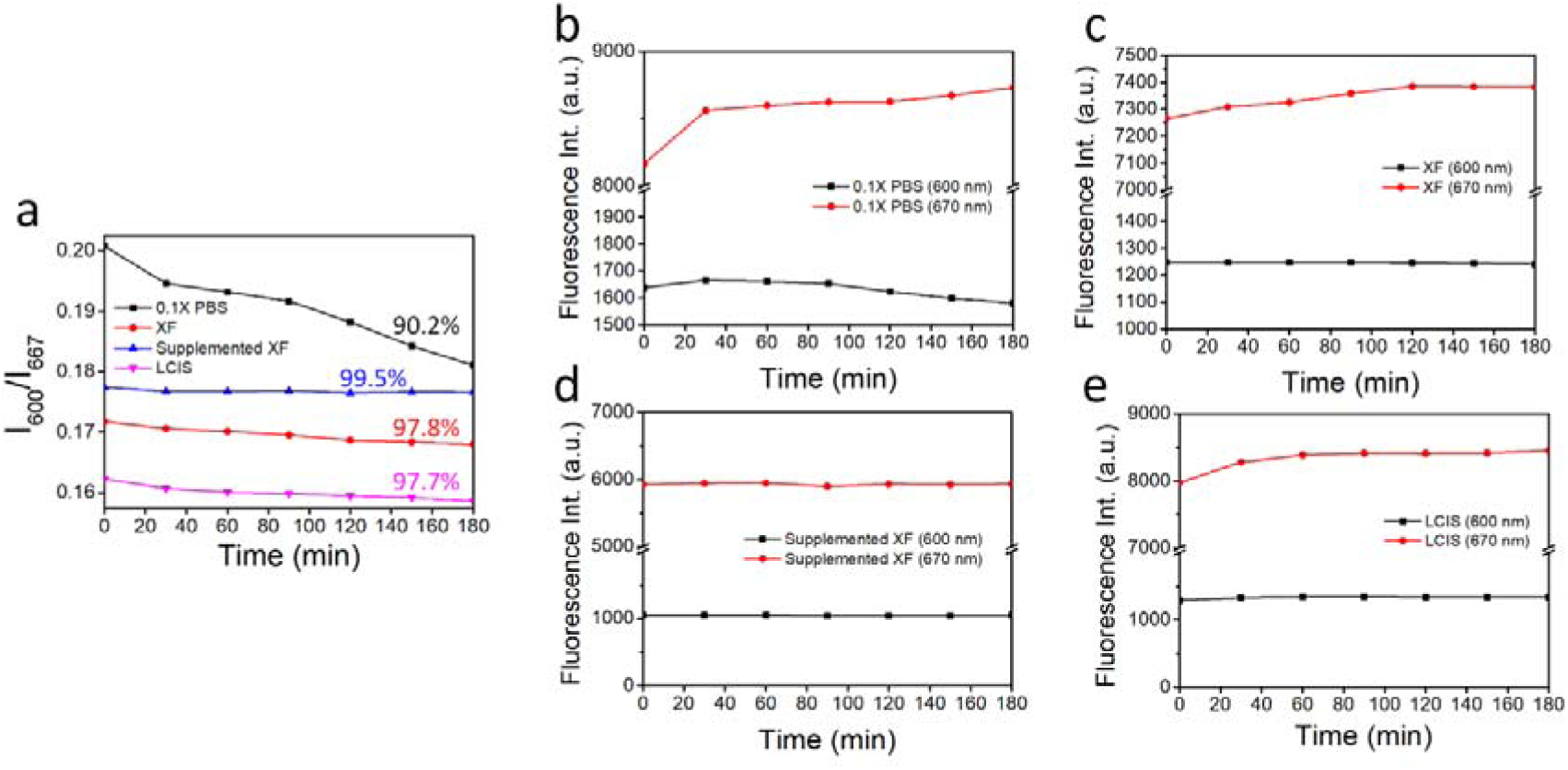
Buffer selection for long-term pH monitoring (37°C, 0% humidity, 0% CO_2_) To accurately measure extracellular pH in tiny droplets, buffer stability under a long-term observation condition has been assessed using 0.1X PBS, Seahorse XF RPMI medium (XF), supplemented XF RPMI medium, and live cell imaging solution (LCIS). XF RPMI medium is supplemented by 2 mM of GlutaMax, 10 mM of glucose, and 1 mM of sodium pyruvate according to the reference [6] 50 μM of SNARF-4F was prepared in four different buffer conditions and was loaded into the Secure-Seal^TM^ hybridization chamber for observation under the microscope. Imaging was performed at 600 nm and 667 nm for 3 hours at 5 min intervals. After imaging, fluorescence intensities at each wavelength were compared. (a) The ratio of fluorescence intensities at 600 nm (I_600_) and 667 nm (I_667_) of four different buffer conditions. Fluorescence intensities at 600 nm and 667 nm of (b) 0.1X PBS, (c) XF, (d) supplemented XF medium, and (e) LCIS.

**Video S1. Cell tracking in droplets by particle tracking analysis program.** Hyperglycolytic (HG) cells are detected by the GFP channel, and untreated (UT; normal glycolysis) cells are detected by the DAPI channel. Position 1 shows three droplets containing HG multiple cells, HG single cell, and UT single cell in order from left to right. Position 2 shows three droplets containing UT multiple cells, UT single cell, and no cell in order from left to right.

**Figure S8.**
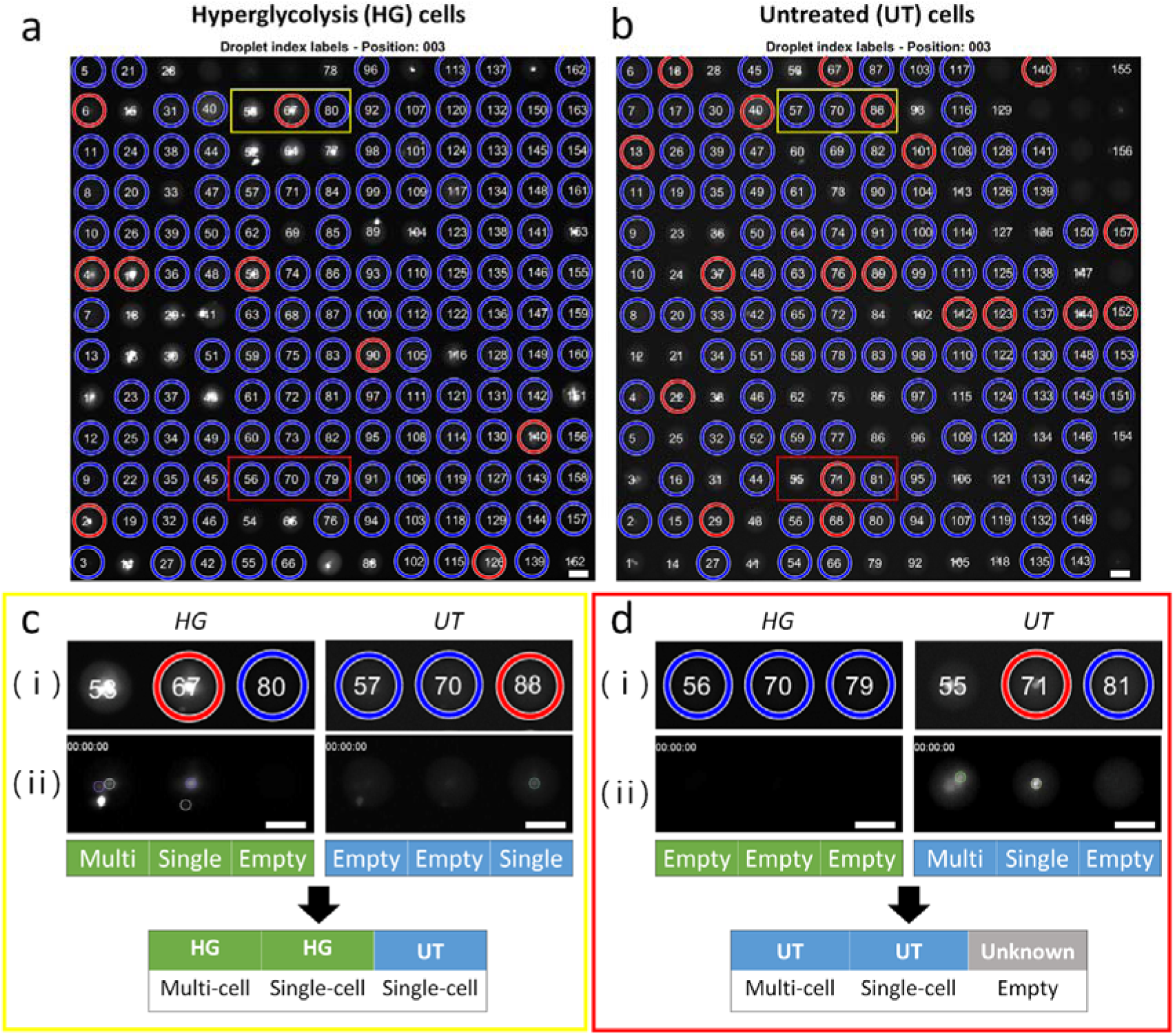
HG and UT droplet discrimination method details. Droplet indexing results are obtained from MATLAB: (a) Hyperglycolytic (HG) cells are detected at the GFP channel, and (b) untreated (UT) cells are detected by the DAPI channel. Droplets containing a single cell are indicated as a red circle, empty droplets are indicated as blue circles, and droplets containing multiple cells have no marks. For examples of pH monitoring performance, three droplets in the yellow boxes and red boxes in both images are magnified in (c) and (d), respectively. (c) (L) Magnified droplet indexing images of HG cells and UT cells, and (L) the images of detected cells by the Particle Tracker are displayed. Detected single cells are outlined in the representative image at the beginning of imaging (0h). The type of droplet (Hyperglycolytic cells vs. Untreated cells; HG vs. UT) and number of cells (Single vs. Multi vs. Empty) are indicated in the table. (d) (L) Magnified droplet indexing images of HG cells and UT cells in red boxes. (L) Images of the detected cells by the Particle Tracker are displayed below the droplet indexing images. Cells are outlined in the image at the beginning of imaging (0h). The scale bar is 50 μm. To enhance the reader’s understanding, we show how we distinguish droplets containing HG cells and UT cells that are randomly immobilized in a microwell array. By comparing droplet indexing results of GFP and DAPI images, which are corresponding to HG cells and UT cells. We can know that three droplets in a yellow box are not HG multi cell droplet, HG single cell droplet, and empty droplet and are HG multi cell droplet, HG single cell droplet, and UT single cell droplet. In the red box, only UT cells are observed. Three droplets are UT multi cell droplet, UT single cell droplet, and empty droplet. Based on these results, we determine that this technique can monitor single cell acidification with diverse glycolytic metabolism.

**Figure S9.**
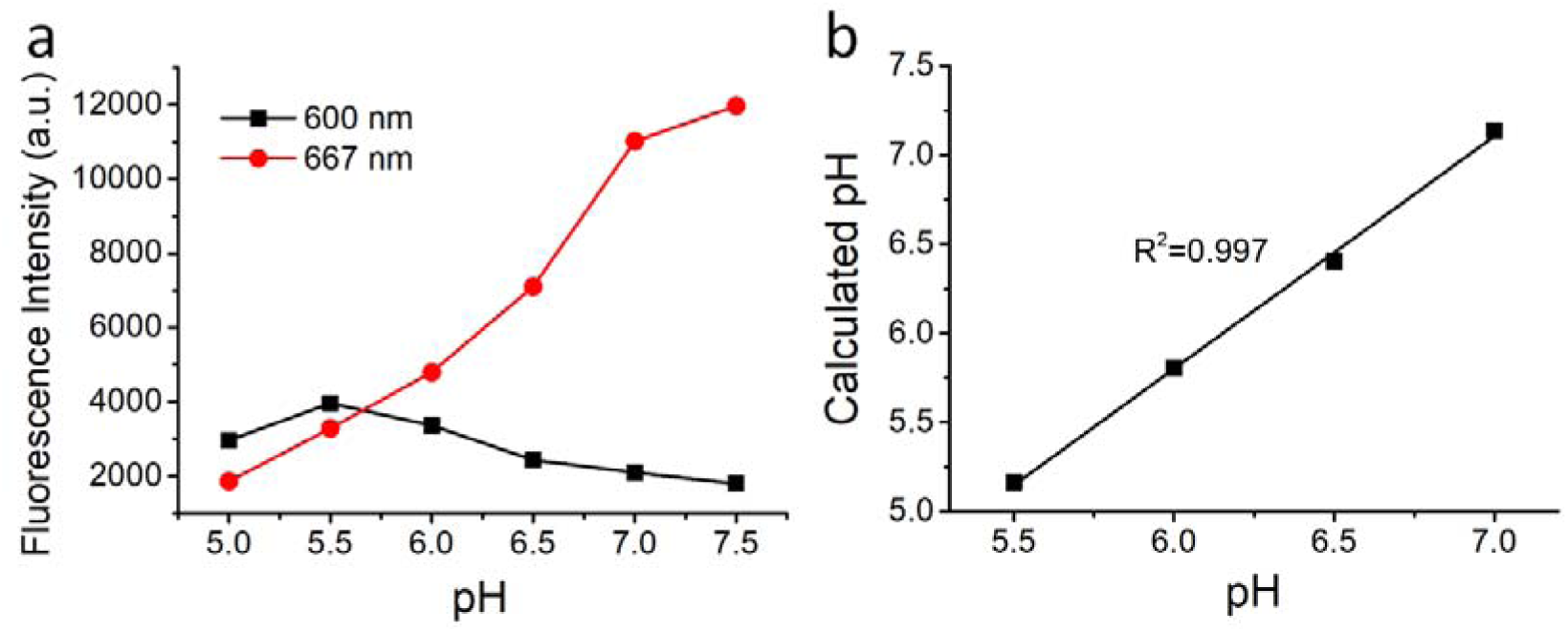
Calibration curve of 50 μM of SNARF-4F in XF unbuffered solution with supplements obtained by a fluorescence microscope. (a) Fluorescence intensities at 600 nm and 667 nm in the pH range from 5 to 7.5. (b) A calibration curve calculated by equation (1) provided by a manufacturer. Calculated pH corresponds to the actual buffer pH (R^2^=0.997). 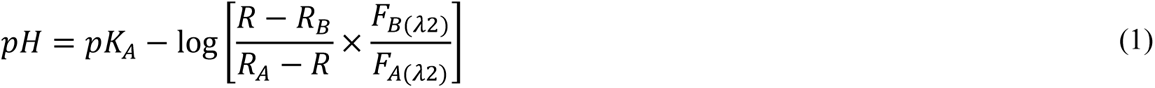 The pH-dependent shift of SNARF-4F allows calibration of the pH response using the ratio of fluorescence intensities measured at the two emission wavelengths. R is the ratio / of fluorescence intensities (F) measured at two emissions and and fixed excitation at 555 nm. R_A_ and R_B_ represent the limiting values at the acidic and basic endpoints of the titration respectively. / represents the normalization factor. pK_A_ of SNARF-4F is 6.4.

## Notes

### Competing Interest Statement

The authors have declared no competing interest.

### Summary of Updates

Could you please add Acknowledgements, Conflicts of interest, and Author contribution as below, after removing the present Acknowledgements? Thank you very much. Acknowledgements Research reported in this publication was supported by the National Human Genome Research Institute of the National Institutes of Health under Award Number RM1HG010023. The content is solely the responsibility of the authors and does not necessarily represent the official views of the National Institutes of Health. E.A.L.P. is funded through the University of Pennsylvania Fontaine Society. Conflicts of Interest The authors declare no conflict of interest. Author Contributions H.J., S.H.H., D.L., and J.K. designed the study. H.J. implemented experiments, analyzed the data, and wrote the manuscript. E.A.L.P. and B.L. created the image analysis pipeline. D.K. wrote the image file sorting script. The other authors assisted with the experiments and discussed the results. All authors have read and approved the final version of the manuscript.

